# Adducins regulate morphology and fate of neural progenitors during neocortical neurogenesis

**DOI:** 10.1101/2024.11.08.622634

**Authors:** Chiara Ossola, Nikola Cokorac, Stefania Faletti, Emanuele Capra, Ilaria Bertani, Chiara Ambrosini, Giovanni Faga, Nereo Kalebic

## Abstract

The evolutionary expansion of the mammalian neocortex is mediated by an increase in the proliferative capacity of neural progenitor cells. However, the molecular machinery controlling the proliferation of apical and basal progenitors during neocortical development is still poorly understood. Here we show that the three actin-associated morpho-regulatory adducins (ADD1-3), underlie abundance of basal progenitors in developing mouse and ferret neocortex *in vivo* and in human cortical organoids. Over expression of adducins in embryonic mouse neocortex increases the number of protrusions of basal progenitors, leading to an increase in their proliferative capacity and neuronal output. Conversely, knock-out of ADD1 in human cortical organoids, which also leads to down-regulation of other adducins, results in reduced proliferation of progenitors and aberrant neurogenesis. Hence, we show that adducins underlie proliferation and fate of neural progenitors, which are key cellular features underlying progression of mammalian neurogenesis.

## Introduction

The evolutionary expansion of human neocortex is widely considered to underlie our advanced cognitive abilities. The current view is that increased proliferative capacity of neural progenitor cells (NPCs) provides foundation of such expansion (Del-Valle-Anton and Borrell, 2022; Kalebic and Huttner, 2020; Lui et al., 2011; Molnar et al., 2019; Rakic, 2009; Sousa et al., 2017; Sun and Hevner, 2014). Namely, human NPCs are thought to undergo multiple proliferative divisions, generating more NPC, which finally results with increased neuronal production and a larger neocortex. NPCs, however, are not uniform and, based on their cell biological features, site of mitosis and gene expression we can distinct two main types: Apical progenitors (APs) and Basal progenitors (BPs) (Taverna et al., 2014; Uzquiano et al., 2018). Whereas APs remain attached to the ventricular surface throughout their cell cycle, BPs delaminate and undergo mitosis at a more basal position, within the subventricular zone (SVZ). Considering the limited space around the ventricle, the ability of BPs to undergo mitosis at a basal position is thought to provide a key foundation for increasing the NPC proliferative capacity and hence the neocortical growth. Interestingly, APs with a continuous contact with the ventricular surface, are highly proliferative cells in all mammals, but BPs seem to be more abundant and more proliferative in species with an expanded neocortex (such as human, macaque or ferret) than in species with a smaller neocortex (such as mouse) (Kalebic and Huttner, 2020; Lui et al., 2011; Namba and Huttner, 2017).

At early stages of neurogenesis APs undergo vertical symmetric division to expand their pool (Taverna et al., 2014). Later, they start dividing asymmetrically either with an oblique or horizontal division angle which promotes generation of delaminated progeny, most often a BP and only rarely a differentiated neuron (Lancaster and Knoblich, 2012; Matsuzaki and Shitamukai, 2015; Penisson et al., 2019). BPs delaminate and migrate into SVZ where they undergo a cell division. A hallmark of proliferative BPs is to sustain those cell divisions in the environment away of the pro-proliferative signals from the ventricle. In mouse such ability is quite low and BPs typically divide only once. In humans, however, this ability is far greater and BPs undergo multiple proliferative divisions. This is enabled through both environmental and cell intrinsic factors. Among the former, extracellular matrix components, growth factors and interactions through cell-cell contacts (Kalebic et al., 2017; Penisson et al., 2019) shape a proliferative niche in the SVZ, thus sustaining BPs proliferation. Instead, an intrinsic feature underlying BP proliferation is their cell morphology. In species with an expanded neocortex, BPs contain more cellular protrusions (Kalebic et al., 2019), which enable them to sense and respond more efficiently to pro-proliferative environmental cues, which sustain their cell cycle re-entry (Kalebic and Huttner, 2020).

However, several important questions remain unanswered. It is still poorly understood how these processes are regulated at the cellular and molecular levels. It is not clear how BP morphology is established and maintained. We have previously identified PALMD, a morphoregulatory molecule that regulates the number of protrusions of human BPs (Kalebic et al., 2019). PALMD is a palmitoylated protein attached to the plasma membrane in a complex with several morphoregulatory proteins, among which adducins take a prominent role. Adducins (ADDs) are a family of membrane skeleton proteins consisting of three members: ADD1, ADD2 and ADD3. They are primarily involved in the assembly of the spectrin-actin cytoskeleton and thus provide physical support to the plasma membrane (Gonzalez-Fernandez et al., 2022; Kiang and Leung, 2018; Matsuoka et al., 2000). Interestingly all three ADDs interact with PALMD in human fetal brain (Kalebic et al., 2019), suggesting their potential role in morphology of human BPs. Furthermore, variants in ADDs have been associated with various neurodevelopmental disorders, including malformations of cortical development, intellectual disability (Qi et al., 2022), inherited cerebral palsy (Kruer et al., 2013) and schizophrenia (Bosia et al., 2016). Overall, these data suggest that ADDs might play an important role during neocortical development.

Here we examined the role of ADDs in mammalian neurogenesis using three different model systems (mouse and ferret *in vivo* and human cortical organoids) and show that this family is both sufficient and required for the correct proliferation of BPs. We found that ADDs regulate the number of protrusions of BPs and orientation of the mitotic spindle in APs, thus controlling proliferation and cell fate of NPCs. Therefore, ADDs underlie key progenitor features enabling the evolutionary expansion of the neocortex.

## Results

### Adducins (ADDs) are required for the correct abundance of basal progenitors in ferret embryonic neocortex

ADDs operate as heterodimers and tetramers of alpha-adducin (ADD1) with either beta-adducin (ADD2) or gamma-adducin (ADD3) (Joshi et al., 1991; Matsuoka et al., 2000). Whereas ADD1 and ADD3 are ubiquitously expressed, although enriched in the brain, ADD2 is restricted to brain and hematopoietic tissues (Bennett et al., 1988; Dong et al., 1995; Kiang and Leung, 2018). Given their potential role in NPC morphology and thereby cell proliferation and neurogenesis, we first sought to examine the histological and cellular expression pattern of ADDs in developing mammalian neocortex. At the mRNA level (Fietz et al., 2012; Florio et al., 2015) ADD1 is expressed ubiquitously in mouse and human developing neocortex, whereas ADD2 is enriched in neurons and ADD3 in NPCs of both species (Figure S1A). We examined the expression of the ADD proteins by immunofluorescence in embryonic mouse and ferret neocortices and human cortical organoids (Figure S1B-D). Similarly to the mRNA expression, ADD1 is present throughout the embryonic cortical wall, but with a marked increase in mitotic APs. Instead, ADD2 is enriched in cortical plate and fiber layers containing neuronal cell bodies and processes, and ADD3 is prominent in the germinal zones containing NPCs. Together this shows that all three ADDs are present in the developing neocortices of various mammals.

Given this expression pattern we sought to knock-out (KO) all three ADDs in the embryonic mammalian neocortex *in vivo* to examine their potential link to BPs biology. We choose to perform the triple KO in the ferret, as this model organism faithfully recapitulates multiple key features of the human neocortex development, including expanded subventricular zone, extended period of forebrain development and abundance and morphological heterogeneity of BPs (Gilardi and Kalebic, 2021). *Via in utero* electroporation we performed a CRISPR/Cas9-mediated triple disruption of gene expression in the embryonic day (E) 33 ferret brains, which corresponds to the onset of the outer SVZ (Martinez-Martinez et al., 2016) and confirmed reduced expression of ADDs four days later (Figure S2A-D). Upon triple KO we detected a reduction in Sox2+ BPs in both inner and outer SVZ by immunohistological analysis (Figure 1). These effects were specific to BPs as the abundance of Sox2+ progenitors was not affected in VZ or IZ (Figure S2E, F).

**Figure 1.**
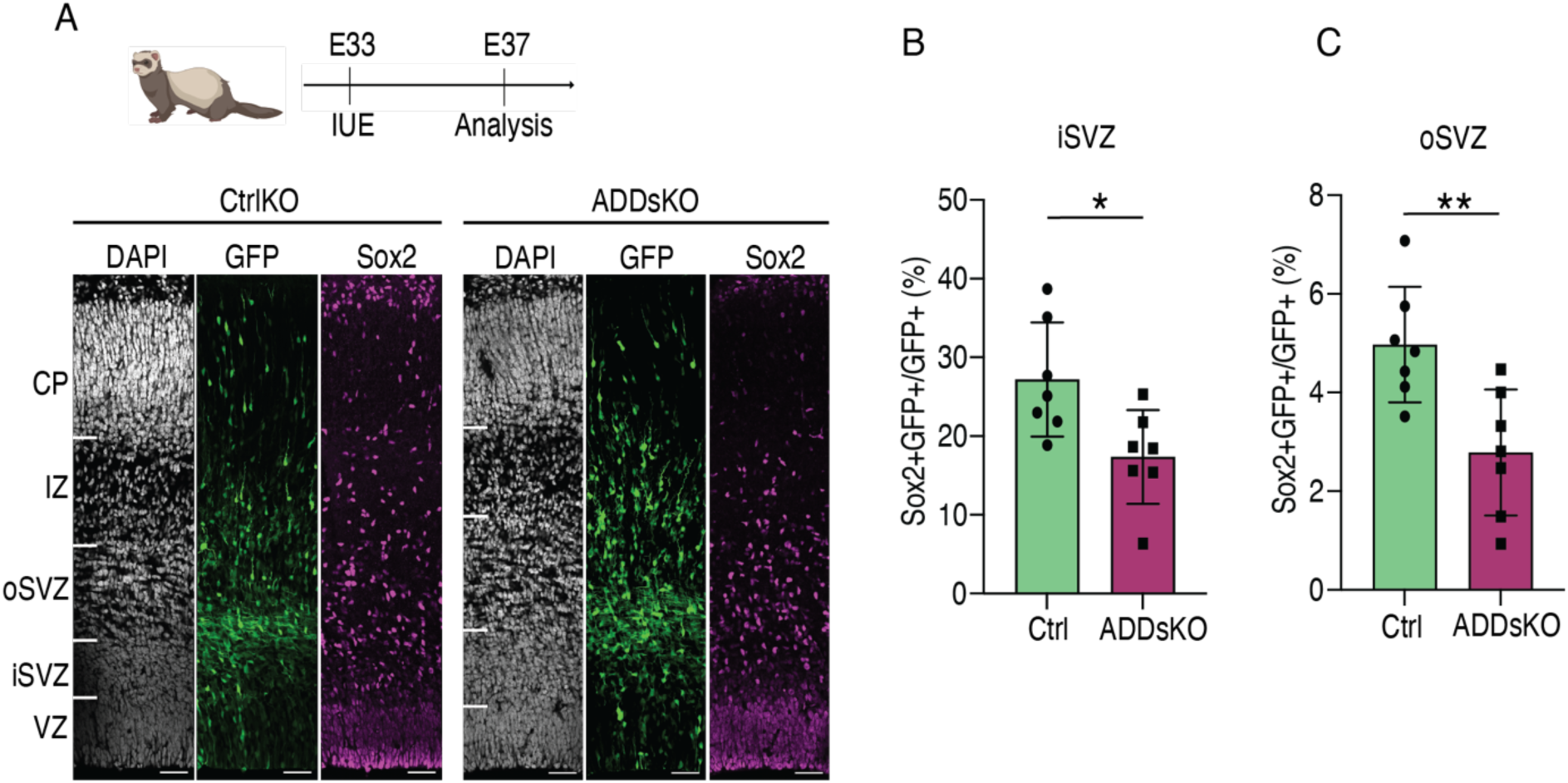
ADDs are necessary for BPs abundance in ferret developing neocortex. ADDs triple KO decreases Sox2+ BPs in both inner and outer SVZ (iSVZ, oSVZ). ADDs KO guides and Cas9 protein were electroporated at E33 along with GFP overexpressing plasmid and embryos were analyzed at E37. (A) DAPI staining and IF for GFP and Sox2 of E37 ferret neocortex for KO (right) and control (left). Scale bar 50 µm. (B, C) ADDs KO significantly decreases Sox2+ BPs in the iSVZ (B) and OSVZ (C). N=7control +7 KO embryos from 3 litters. **p < 0.01, *p < 0.05; Student’s *t* test.

These data suggest that ADDs are required for the correct abundance of BPs in ferret. We hence sought to examine the ability of ADDs to regulate mammalian neurogenesis, using a simpler *in vivo* model system, the embryonic mouse brain.

### Adducins promote NPC abundance in embryonic mouse neocortex

We subcloned each of the three ADDs in pCAGGS vectors (Figure S3A), and over expressed (OE) them, along with the GFP marker, in the embryonic mouse brain at E13 *via in utero* electroporation (Figure S3B-D). This is a developmental stage in which mouse BPs are generated from APs (Arai et al., 2011; Florio et al., 2015) and hence comparable to the approach in ferret (Figure 1).

Given that ADDs operate as heterodimers with ADD1 as an obligatory partner (Matsuoka et al., 2000), we first investigated the effects of ADD1 co-electroporated with either ADD2 or ADD3. Two days after electroporation we examined the abundance of Sox2+ BPs in the mouse SVZ (Figure 2A, D) and detected a strong increase upon co-overexpression (co-OE) of ADD1 with either ADD2 or ADD3 (Figure 2C, F). Conversely, we did not observe any effect on Sox2+ APs in mouse VZ (Figure 2B, E).

**Figure 2.**
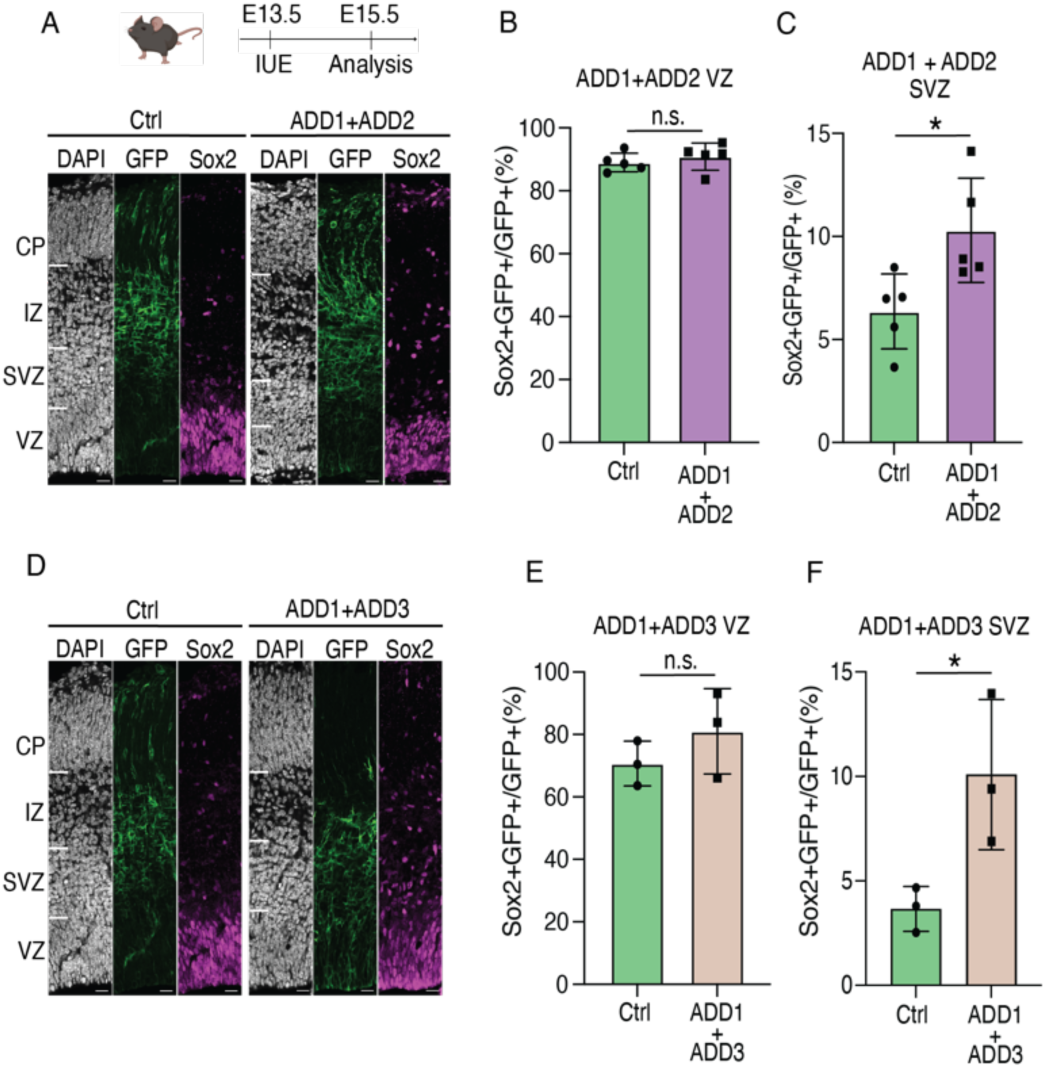
ADDs are sufficient for BPs abundance in mouse developing neocortex. Co-overexpression (co-OE) of ADD1 and ADD2 or ADD1 and ADD3, increase Sox2+ BPs in mouse SVZ, but not Sox2+ APs in VZ. ADDs were electroporated along with LynGFP overexpressing plasmid (revealing cell membrane) at E13.5 in mouse embryonic neocortex and embryos were analyzed at E15.5. (A, D) DAPI staining and IF for GFP and Sox2 of E15.5 mouse neocortex for control (left) and ADD1 and ADD2 co-OE (right, A) or ADD1 and ADD3 co-OE (right, D). Scale bars, 20 µm. (B, C, E, F) Co-OE of ADD1 and ADD2 (B, C) or ADD1 and ADD3 (E, F) significantly increase Sox2+ BPs in the mouse E15.5 SVZ (C, F), but does not a]ect abundance of APs in VZ (B, E). N=5 control and 5 OE embryos (B, C) or 3 control and 3 OE embryos (E, F) from 3 di]erent litters. *p < 0.05, n.s. not statistically significant; Student’s *t* test.

We next examined the effects of individual ADDs to the abundance of Sox2+ NPCs in mouse embryonic neocortex (Figure S3E, H, K). Our analysis showed that only OE of ADD1 resulted in an increase in abundance of Sox2+ APs in the VZ (Figure S3F, I, L), whereas OE of both ADD1 and ADD2 individually, but not ADD3 led to an increase in abundance of Sox2+ BPs in the SVZ (Figure S3G, J, M). Together, these data show that either ADD1 alone or in combination with ADD2 are key adducins affecting the abundance of mouse NPCs, so henceforth we focused on them.

### ADD1 and ADD2 operate together to promote proliferation of mouse BPs

Whereas Sox2+ BPs are the key type of BPs in human and ferret, in mouse the most dominant population are Tbr2+ basal intermediate progenitors (bIPs) (Arai et al., 2011; Taverna et al., 2014). We hence examined the potential effects of the OE of ADD1 alone and in combination with ADD2 to mouse bIPs, however, our immunofluorescence analysis revealed no effect on Tbr2+ bIPs in either case (Figure S4).

The observed increase in Sox2+ BP could be due to increased delamination of Sox2+ APs or due to increased proliferation of BPs. Considering that we haven’t seen any depletion of Sox2+ APs in the VZ (Figure 2B, E), and in fact observed a mild increase in their abundance upon ADD1 OE (Figure S3F), we focused on the latter scenario. We performed a cell cycle re-entry assay by injecting EdU *in vivo* at E14.5, i.e. 24 h after the *in utero* electroporation. Given that mouse BPs have a cell cycle of around 20 h (Arai et al., 2011) we analyzed the embryos at E15.5, i.e. 24 h after the injection. We quantified the proportion of Ki67+ EdU+ cells, i.e. the cells that accumulated EdU in the previous cycle and have since re-entered the next cell cycle. Our analysis showed an increase in such cells upon co-OE of ADD1 with ADD2 (Figure 3A, C), indicating an increase in the BP proliferative capacity. Furthermore, we observed an increase in the abundance of Ki67+ cells in SVZ (Figure 3B), which agrees with the previously observed increase in Sox2+ BP (Figure 2C). The increase in Ki67+ EdU+ cells suggests that such increased abundance is due to an increase in the proliferative capacity of BPs upon co-OE of ADD1 and ADD2. In contrast, APs in VZ did not show increased cell cycle re-entry upon co-OE of ADD1 and ADD2 (Figure S5B), although a mild increase in the abundance of Ki67+ cells was detected in the VZ (Figure S5A).

**Figure 3.**
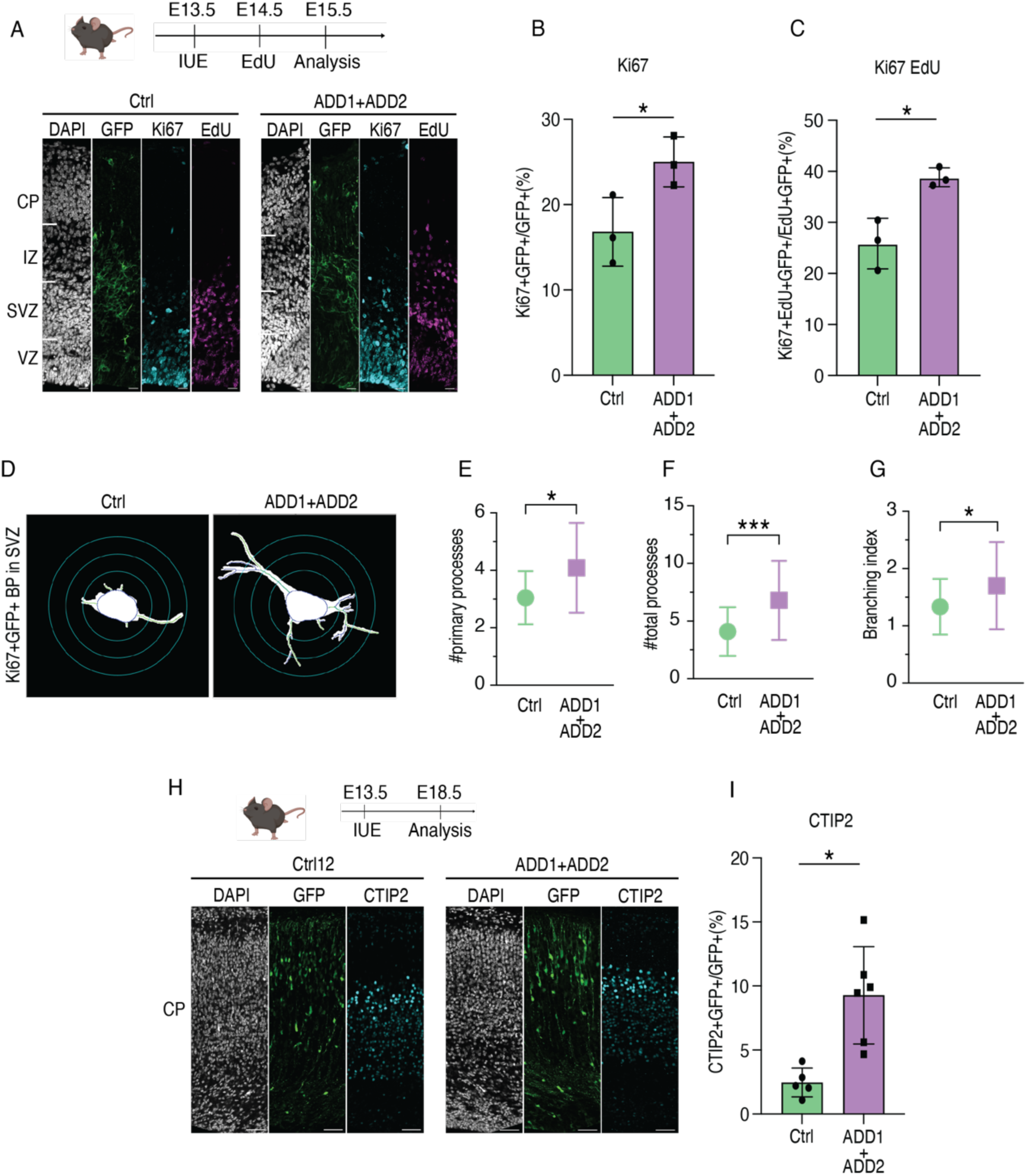
ADD1 and ADD2 co-OE in mouse embryonic neocortex increases BP proliferation and abundance of cellular protrusions. (A-C) ADD1 and ADD2 co-OE increases BPs proliferative capacity and cell cycle re-entry. ADDs were electroporated along with LynGFP overexpressing plasmid at E13.5 in mouse embryonic neocortex, EdU was administered by an intraperitoneal injection at E14.5 and embryos were analyzed at E15.5. (A) DAPI staining and IF for GFP, Ki67 and EdU of E15.5 mouse neocortex for control (left) and ADD1 and ADD2 co-OE (right). Scale bar 20 µm. (B, C) ADD1 and ADD2 co-OE increases abundance of actively proliferating (Ki67+) BPs (B) and the cell cycle re-entry frequency (C) in mouse E15.5 SVZ. N=3 control and 3 co-OE embryos from 3 di]erent litters. *p < 0.05; Student’s *t* test. (D-G) ADD1 and ADD2 co-OE increases the abundance of BP protrusions in mouse developing neocortex. ADDs were electroporated along with GFP overexpressing plasmid at E13.5 in mouse embryonic neocortex and analyzed at E15.5. (D) Segmentation masks of Ki67+GFP+ BPs in mouse E15.5 SVZ of control (left) and ADD1 and ADD2 co-OE (right). (E, G) ADD1 and ADD2 co-OE increases the number of BPs primary (E) and total (F) processes as well as the branching index (G). N=23 control and 34 co-OE cells from 3 di]erent litters. ***p < 0.001; *p < 0.05; Mann-Whitney *u*-test (E, F); Welch’s t test (G). (H-I) ADD1 and ADD2 co-OE increases neuronal generation in mouse developing neocortex. ADDs were electroporated along with GFP overexpressing plasmid at E13.5 in mouse embryonic neocortex and embryos were analyzed at day E18.5. (H) DAPI staining and IF for GFP and CTIP2 of E18.5 mouse neocortex for control (left) and ADD1 and ADD2 co-OE (right). Scale bar 50 µm. (I) ADD1 and ADD2 co-OE significantly increases abundance of CTIP2+ neurons in mouse E18.5 cortical plate. N=6 control and 6 co-OE embryos from 3 di]erent litters. *p < 0.05; Student’s *t* test.

Importantly, to distinguish if ADD1 alone was sufficient for promoting proliferation in SVZ, or its partner ADD2 was also required, we performed OE of ADD1 alone and examine the abundance of Ki67+ NPCs and their proliferative capacity *via* cell cycle re-entry analysis. Remarkably, our data show that ADD1 alone was not able to promote abundance of Ki67+ cells nor their proliferative capacity in either VZ or SVZ (Figure S5C-G).

Taken together, our data suggest that both ADD1 and ADD2 are required for the proliferation of mouse BPs. They further show that ADDs primarily exert their effects on NPCs in the mouse SVZ.

### ADD1 and ADD2 promote growth of BP protrusions and mouse neurogenesis

Given that ADDs are morpho-regulatory proteins and that cell morphology, namely the number of protrusions BPs grow, has been linked to their proliferative capacity (Kalebic et al., 2019; Kalebic and Huttner, 2020), we next examined the morphology of Ki67+ BPs upon OE of ADD1 and ADD2, following an established pipeline (Kalebic et al., 2019). BPs co-overexpressing ADD1 and ADD2 exhibited more protrusions and a more branched morphology compared to control BPs (Figure 3D-G). Of note, OE of ADD1 alone also led to an increase in the number of BP cell protrusions (Figure S5H-K). This was likely not translated to an increase in the BP proliferative capacity, as those protrusions were smaller and with shorter branches compared to the BPs overexpressing both ADD1 and ADD2 (Figure S5L).

Mouse BPs at this stage of mouse neurodevelopment are neurogenic, so to understand the consequences of the increased abundance and proliferative capacity of mouse BPs we sought to examine the abundance of generated neurons. To this end, we co-electroporated mouse embryos with ADD1 and ADD2 at E13 and examined the cortices at E18 (Figure S6A, B). We detected a marked increase in the abundance of CTIP2+ (but not SATB2+) neurons upon ADDs OE compared to the control embryos (Figure 3H, I and S6C, D). Since this effect was absent upon OE of ADD1 alone (Figure S6E-G), it is likely that it is due to the increase in BPs proliferative capacity (Figure 3C). This suggests that the effects of ADDs OE on mouse neural progenitors elicit an increase in mouse neurogenesis.

Together these data show that ADDs increase the number of BP processes and, in agreement with the hypothesis that BP morphology and BP proliferation are closely linked, they also promote the proliferative capacity of those cells which leads to an increase in neuronal production.

### ADD1 is required for progenitor proliferation in human cortical organoids

We next examined whether ADDs are required for human neocortical neurogenesis. To this end, we employed sliced human cortical organoids (Qian et al., 2020) which have been previously shown to sustain neurogenesis, organoid growth and SVZ expansion over a long time period and have been used to study proliferation of human BPs (Cubillos et al., 2024). Importantly, they also express all three ADDs (Figure S1).

Although in the mouse OE studies (Figures 2 and 3), we observed key effects upon co-OE of ADD1 with ADD2, for the loss of function studies in human cortical organoids we generated only ADD1 KO, since ADD1 is known to be the limiting subunit in functional tetramer formation and in its absence no ADDs complex is formed (Robledo et al., 2008). We hence generated three different ADD1 KO lines in H9 human embryonic stem cell background targeting *ADD1* (Figure S7) and subsequently derived cortical organoids, which we examined at two developmental stages, day (D) 45 and D75, which correspond to pre-SVZ and to the peak of SVZ stages, respectively (Figure 4A). We confirmed the depletion of all three ADD proteins in the KO line via immunoblot (Figure S8A). The absence of ADD2 and ADD3 proteins, although their mRNA was present (Figure S8B) agrees with previous reports analyzing ADD1 KO mice (Robledo et al., 2008) and further corroborates that when ADD1 is not present, no ADD complexes forms at the spectrin-actin junctions.

**Figure 4.**
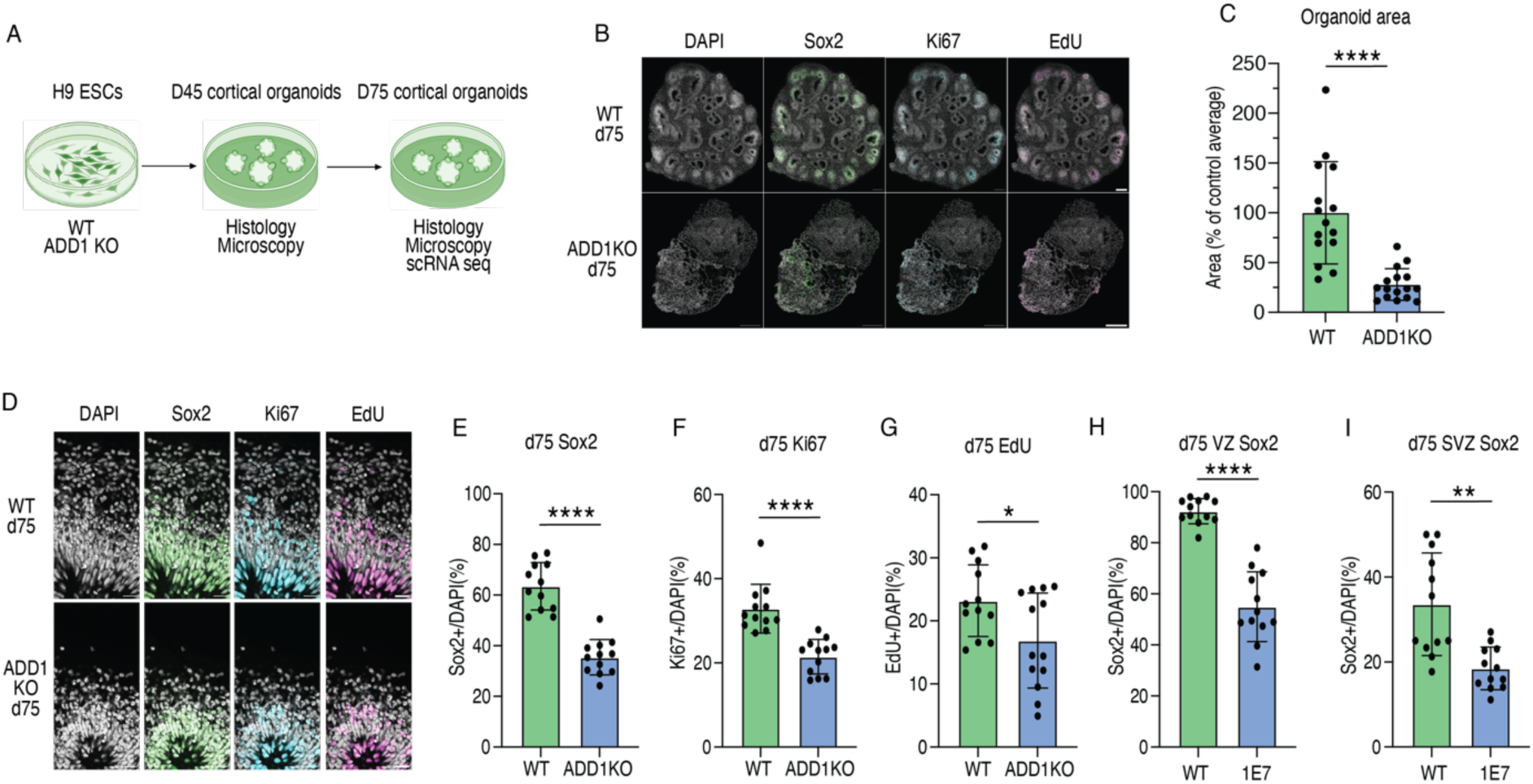
ADD1 KO in human cortical organoids reduces NPC proliferation. WT and 3 di]erent clones of ADD1KO H9 PSCs (see Figure S7) were di]erentiated into cortical organoids (Qian et al., 2020) and analyzed at day 45 (see Figure S8) and day 75 (this figure and Figure 5). (A) Experimental plan. (B, D) DAPI staining and IF for Sox2, Ki67 and EdU of WT (upper) and 1E7 clone of the ADD1 KO (down) at day 75 cortical organoids. (B) overview images; (D) single lumen (ventricle). Scale bars, 100 µm (B), 20 µm (D). (C) ADD1 KO significantly reduces organoids size at day 75. N=15 WT and 15 KO organoids from from 3 di]erent di]erentiation batches. ****p < 0.0001, Welch’s t test. (E-I) ADD1 KO significantly reduces abundance of SOX2+ NPCs (E), Ki67+ cycling cells (F) and EdU+ S-phase cells (G) in the germinal zones around the organoid lumens at day 75. Among the SOX2+ NPCs, both APs in the VZ (H) and BPs in the SVZ (I) are reduced. N= 12 WT and 12-13 KO organoid lumens from 3 di]erent di]erentiation batches. ****p < 0.0001, **p < 0.01, *p < 0.05; Welch’s *t* test (E, G-I), Kolmogorov-Smirnov test (F).

We performed an immunohistological analysis of neural progenitor proliferation at D45 for all three KO lines (Figure S8C) and detected a marked reduction in the abundance of SOX2+ and Ki67+ NPCs as well as EdU+ proliferative cells (Figure S8D-F). Since all three ADD1 KO clones showed similar magnitudes of effect, we performed analyses at later differentiation stage (D75) on one clone (1E7). From three independent differentiations of the clone 1E7 we observed a strong four-fold reduction in the organoid size at D75 compared to WT controls (Figure 4B, C) accompanied by generally aberrant organoid architecture. Immunohistological analysis of D75 organoids (Figure 4D) showed a reduction in progenitors abundance as observed by SOX2 and Ki67 markers(Figure 4E, F) and in proliferation as observed by EdU positivity (Figure 4G). Notably, the abundance of SOX2+ NPC was strongly reduced in both VZ and SVZ (Figure 4H, I), suggesting an effect on both APs and BPs.

In summary, human cortical organoids lacking ADD1 showed reduced growth due to reduced NPC proliferation. We next sought to (1) gain a mechanistic insight into the cellular alterations leading to reduced proliferation and to (2) examine the consequences of the ADD1 KO on the progression of human neurogenesis.

### ADD1 is required for correct progression of human neurogenesis

To gain insight into the transcriptomic signature of the cellular alterations underlying reduced NPC proliferation upon ADD1 KO, we performed single cell RNA sequencing (scRNA-seq) of D75 cortical organoids. We sequenced 33 WT and 36 ADD1 KO organoids coming from 4 (WT) and 3 (ADD1 KO) differentiations and obtained ∼49k cells that passed all the quality control streps. Leiden clustering revealed 5 principal clusters pertinent to human mid neurogenesis (Cycling progenitors, G1 Radial glia (RG), G1 intermediate progenitors (IPs) and 2 neuronal clusters) in addition to a minor cluster of other cells (Figure 5A and see Figure S9A for the markers of individual clusters). Among the neuronal clusters, Neurons 1 were enriched in early-born and mature neurons (TBR1+, CTIP2+), whereas Neurons 2 cluster contained late-born and upper-layer neurons (CUX2+) (Figure S9A). While both WT and ADD1 KO contributed to all clusters we observed unequal relative distribution of cells within the Neurons 2 cluster and we hence performed subclustering of Neurons 2 cluster and identified Neurons 3 subcluster, which showed a marked enrichment in the KO (Figure 5B, C).

**Figure 5.**
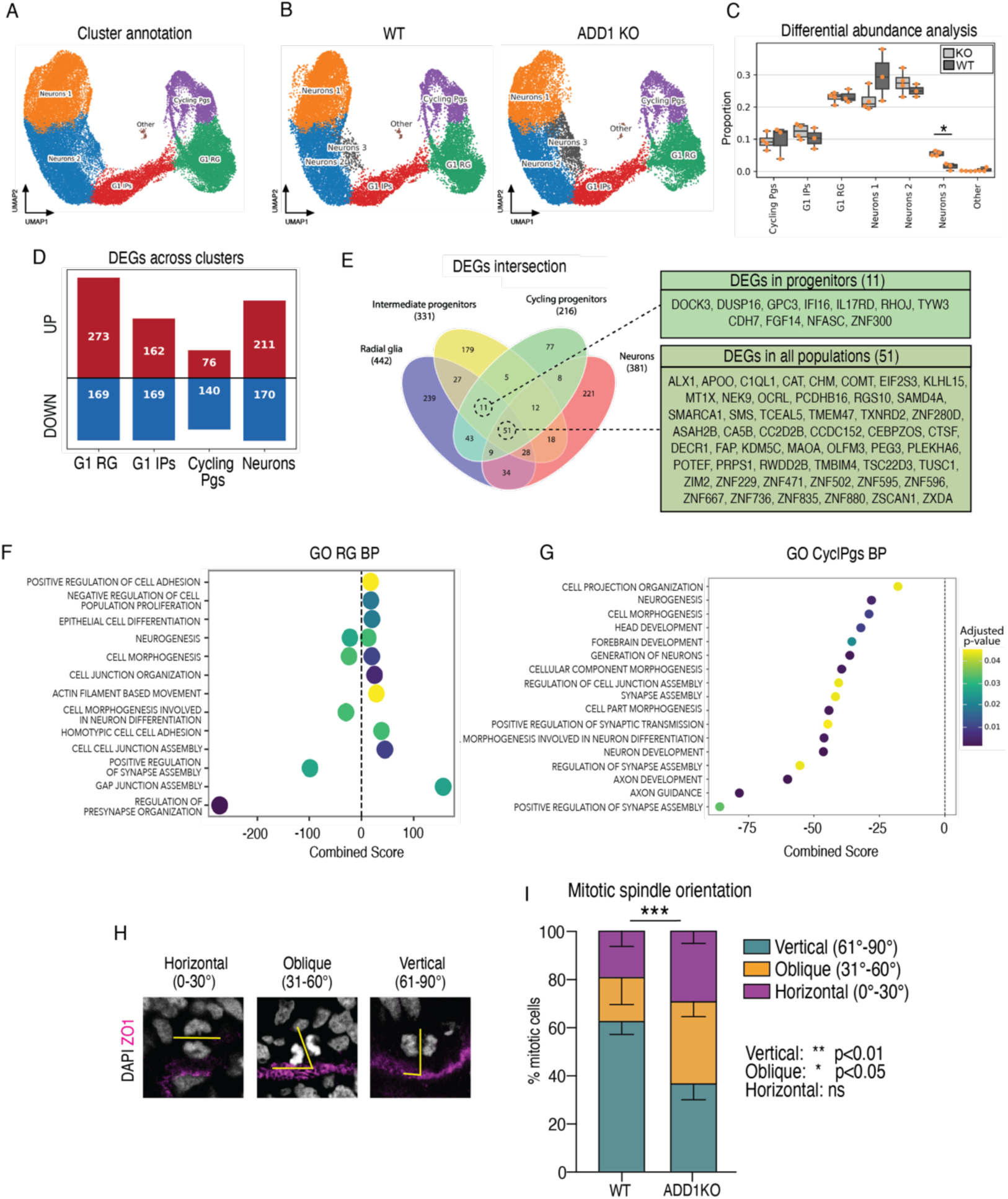
ADD1 is required for correct progression of human neurogenesis and spindle orientation of APs. (A-G) ScRNA-seq analysis of WT and ADD1 KO cortical organoids at day 75. (A) UMAP embedding of scRNA-seq data (entire dataset) with the identified clusters of cells (G1 Radial glia, Cycling progenitors, G1 intermediate progenitors, Neurons 1 and 2). (B) Genotype-based (WT, left; ADD1 KO, right) UMAP embedding of scRNA-seq data showing Neurons 3 subcluster. (C) Di]erential abundance analysis (scCODA) of principal cell clusters showing an enrichment in Neurons 3 subcluster. N = 4 WT and 3 ADD1 KO di]erentiation batches. *p < 0.05. (D) Number of identified di]erentially expressed genes (DEGs) across cell clusters (ADD1 KO vs. WT). (E) Schematic representation of DEGs intersections across clusters, listing the 51 DEGs in all clusters and further 11 DEGs shared by all NPCs. (F) Plot showing selected gene ontology (biological process) terms enriched in radial glia (GO RG BP) in ADD1 KO vs. WT. For the whole list see Figure S10. (G) Plot showing all GO BP enriched terms in cycling progenitors (CycPgs) in ADD1 KO vs. WT. (H) DAPI staining and IF for ZO1 of WT cortical organoids (day 45) showing examples of horizontal, oblique and vertical mitotic spindle orientation. (I) ADD1 KO significantly reduces vertical mitotic division with a relative increase in oblique divisions. N= 76 WT and N=95 KO apical mitoses from 3 di]erent di]erentiation batches. ***p < 0.001, **p < 0.01, *p < 0.05, n.s. statistically not significant; 2-way ANOVA and Bonferroni’s post-hoc test.

Whereas Neurons 3 contained the markers of immature neurons along with the synaptic markers, their expression levels were reduced compared to the other neuronal clusters (Figure S9A). Gene onthology (GO) analysis of genes differentially expressed between Neurons 3 and their paternal cluster (Neurons 2) showed reduced neurogenesis and cellular morphogenesis along with increased response to ER stress (Figure S9B, D). Specifically, they expressed multiple markers of the PERK-ATF4 stress pathway and showed reduced expression of mitochondrial genes (Figure S9C). Furthermore, a pseudotime analysis corroborated that Neurons 3 are less advanced than other neurons on the differentiation trajectory (Figure S9E, F). Single-cell fate mapping (Lange et al., 2022) predicted that the differentiation trajectory leading to Neurons 3 is more likely to initiate in the RG, rather than IP, cluster. Together, these data suggest that ADD1 KO leads to aberrant neurogenesis with an accumulation of immature neurons in a state of cellular stress.

To gain transcriptomic insight into the cellular mechanisms underlying the effects of ADD1 KO we performed a differential gene expression analysis on the entire dataset and detected 962 differentially expressed genes (DEGs) (Figure 5D). Interestingly, 51 genes were differentially expressed in all clusters and further 11 in all NPC clusters, suggesting a general transcriptomic signature of ADD1 KO (Figure 5E). However, the greatest number of DEGs was in the RG cluster (Figure 5D). GO enrichment analysis of DEGs in this cluster showed that genes up-regulated upon ADD1 KO show enrichment for the terms associated with cell adhesion and cell-cell junctions (Figure 5F and S10, S11A, E, F). Instead, the genes down-regulated in this cluster contributed to the enrichment in GO terms such as cell morphogenesis in neuronal differentiation (Figure 5F, S11A). Similarly the genes down-regulated in ADD1 KO cycling progenitors were also enriched in terms related to cell morphogenesis and neurogenesis progression (Figure 5G, S11B, G, H). In contrast, the GO analysis of DEGs in IPCs and neurons (Figure S11C, D) did not yield any significant terms. Together, this suggests that ADD1 KO causes alterations in NPCs, and in particular in RG, that then can lead to altered cell morphology and impaired neurogenesis.

### ADD1 is required for correct spindle orientation of human APs

The aberrant neurogenesis observed upon ADD1 KO (Figure 5B, C) could be due to effects of ADDs directly on neuronal maturation or due to their upstream effects in NPCs, notably RG, which regulate generation of neurons. Although we detected a general signature in all cells lacking ADDs (Figure 5E), our scRNA-seq analysis primarily suggests that the effects are likely due to cellular events in RG (Figure 5F and S9H, S10, 11). The altered fate of NPC progeny (Figure 5B, C, S9H) and the reduced organoid size (Figure 4C) together prompt us to examine the division mode of RG. We hypothesized that the observed phenotypes might be linked to mitotic spindle orientation, which is a key determinant of asymmetric cell fates, influencing thereby generation of BPs, neuronal output and brain size (Lancaster and Knoblich, 2012; Wynshaw-Boris, 2013). Considering that ADD1 has been previously implicated in the spindle assembly and integrity in other systems (Chan et al., 2014; Hsu et al., 2018) we sought to examine the mitoses of D45 cortical organoids. We calculated the angle of division upon immunofluorescence for ZO1, a marker of the luminal (ventricular) surface, and staining for DAPI to reveal the chromatin (Figure 5H). At this stage of the organoid development, WT cells undergo 60% of vertical division generating two APs that remain attached to the ventricle (Figure 5I).

Upon ADD1 KO however we observed a marked decrease in the vertical divisions with a relative increase in oblique ones (Figure 5I). This suggests that upon ADD1 KO there is a depletion of proliferative APs and increased generation of immature delaminated cells which then leads to aberrant neurogenesis.

## Discussion

Our study identifies fundamental roles of actin-binding adducins (ADDs) in neocortical development. The following aspects of the study deserve particular discussion: (1) role of ADDs in morphology of BP; (2) role of ADDs in cell fate determination; along with their implications (3) for the progression of neurogenesis and (4) for the evolutionary expansion of neocortex and neurodevelopmental pathologies.

### ADDs regulate morphology of BPs

We identified regulation of BP morphology as a key role of ADDs in neurogenesis, as they were able to increase the number of BP cellular protrusions and their branching (Figure 3D-G, S5H-L). We have previously shown that the depletion of ADD3 in human fetal neocortex leads to a reduction in the number and length of BP cellular protrusions (Kalebic et al., 2019). Together, this suggests a key role of ADDs in controlling BP morphology.

ADDs are involved in the assembly of the spectrin-actin cytoskeleton by capping the fast-growing end of actin filaments and recruiting additional spectrin subunits thereby preventing the addition or loss of actin subunits (Bennett et al., 1988; Gardner and Bennett, 1987; Kuhlman et al., 1996; Li et al., 1998; Matsuoka et al., 2000; Taylor and Taylor, 1994). We hypothesize that this role in maintain the cytoskeleton stability could be linked with ADDs’ ability to promote cellular protrusions in BPs. In agreement with this, ADDs have been implicated in promoting outgrowth of neurites and consequently synaptic plasticity in neurons (Lou et al., 2013; Matsuoka et al., 1998; Porro et al., 2010; Yaguchi et al., 2017). Furthermore, ADDs were shown to promote growth of cellular protrusions in cells in culture (Chen et al., 2007) and regulate morphology of platelets (Barkalow et al., 2003), suggesting a general role in maintaining cell morphology. We have previously identified PALMD, as one of key morphoregulatory proteins in BPs and considering that ADDs interact with PALMD (Kalebic et al., 2019) it is plausible that these proteins operate together to maintain BP morphology. Given the role of ADDs in stabilizing actin cytoskeleton we speculate that the PALMD-ADDs complex maintains stability of the newly-formed protrusions, although a direct effect on the process growth could also be possible.

The cell morphology of BPs is associated with their proliferative capacity (Kalebic and Huttner, 2020; Kalebic and Namba, 2021). To support BP proliferation away from the pro-proliferative signals of the cerebrospinal fluid, a stem cell niche in the SVZ (and most notably human outer SVZ) provides such signals. Hence growing additional cellular protrusions can enable BPs to receive more pro-proliferative signals and thereby enable them to undergo repetitive proliferative divisions, which in turn increases their abundance (Kalebic and Huttner, 2020). A variety of such extrinsic pro-proliferative signals has been identified over the years, ranging from ECM components, growth factors, morphogens and various molecules mediating cell-cell contact (Ferent et al., 2020).

ADDs operate in heterodimers with ADD1 as an obligatory partner (Joshi et al., 1991; Matsuoka et al., 2000). Interestingly, we observed an increase in BP protrusions both upon OE of ADD1 alone and upon co-OE of ADD1 with ADD2, but we noticed an effect on BP proliferation only upon the co-expression (Figure 3 and S5). Considering that endogenous ADDs are present in the mouse embryonic neocortex at this stage (Figure S1), we hypothesize that the OE of individual ADDs might promote formation of complexes with endogenous mouse proteins, or even increase their expression levels, and therefore lead to subtle effects on NPC abundance. Such effects appear to be the strongest upon OE of ADD1 alone, which is in line with the notion that ADD1 is obligatory partner of the complex. However, the interactions with endogenous proteins were not sufficient to promote proliferation of mouse BPs. Importantly, we detected slightly more protrusions and more distant branches with respect to the cell body upon the co-expression of ADD1 with ADD2 compared to OE of ADD1 alone (Figure S5L). This might indicate that in order to achieve an effect on cell proliferation a certain threshold in the number or span of cellular protrusions is required, which in turn is enabled by the sufficient amount of ADDs. This is a tempting hypothesis as the amount of the cell surface could be linked to the abundance of various receptors that receive the extrinsic pro-proliferative signals. In agreement with this, we have previously shown that the number of BP protrusions increases from mouse to ferret to human together with the increase in the proliferative capacity of these cells (Kalebic et al., 2019). Furthermore, considering that the OE of morphoregulatory PALMD led to accumulation of integrins on newly-induced processes (Kalebic et al., 2019), it would be interesting to examine if OE of ADDs also results in recruitment or upregulation of membrane receptors.

### ADD1 and cell fate determination

We have identified regulation of the mitotic spindle orientation of APs as the earliest effect of ADDs on the human neurogenesis trajectory. The spindle orientation is a key element determining symmetric *vs* asymmetric cells division (Matsuzaki and Shitamukai, 2015; Taverna et al., 2014) and in neurogenesis it is tightly linked to the asymmetric cell fate and generation of BPs and neurons thereby influencing neuronal output and neocortical expansion (Fish et al., 2008; Lancaster and Knoblich, 2012). When the spindle orientation is vertical it leads to generation of two APs that remain attached to the ventricle. Interestingly, in our WT organoids around 60% of spindles has this orientation suggesting that the pool of APs is expanding (Figure 5I). Oblique orientations instead lead to asymmetric division and generation of delaminated and often more differentiated progeny (basal intermediate progenitors or neurons) (Shitamukai et al., 2011). Finally, horizontal division angle is strongly associated with the generation of basal radial glia (bRG) in humans (LaMonica et al., 2013). Upon ADD1 KO, which leads to depletion of all ADDs, we have detected an increase in oblique division with a relative decrease in vertical ones, but no effect on horizontal divisions. We interpret that this leads to a depletion of APs and reduced generation of bRG, which is in accordance with our data showing a decrease of SOX2+ cells in both VZ and SVZ (Figure 4H, I).

Actin and myosin play an important role in the organization and function of the mitotic spindle (di Pietro et al., 2016; Kunda and Baum, 2009; Rosenblatt et al., 2004; Woolner et al., 2008). In this context ADD1 has been previously implicated in the assembly and integrity of the mitotic spindle, through interactions with myosin-X and TPX2 (Chan et al., 2014; Hsu et al., 2018). ADD1 knock down has been shown to lead to distorted and elongated spindles, defective spindle poles and centriole splitting (Chan et al., 2014; Hsu et al., 2018). In the future it would be interesting to examine the integrity of the mitotic spindle in ADD1 KO cortical organoids to identify the specific role of ADDs in this process. Finally, alterations in spindle orientation during neurodevelopment often lead to cortical malformations and most notably, microcephaly (Gilmore and Walsh, 2013; Juric-Sekhar and Hevner, 2019; Ossola and Kalebic, 2021). This is typically due to depletion of the NPC pool lining the ventricles and premature neuronal differentiation, which ultimately results in the reduced brain size. It is hence interesting to note that upon the depletion of ADD1, we detected a remarkable reduction in the size of human cortical organoids (Figure 4C).

### ADDs and progression of neurogenesis

As suggested above, both effects on APs and BPs can ultimately lead to consequences on neuronal output. We show that OE of ADDs leads to increased neuronal production in mouse, whereas depletion of ADDs resulted in an aberrant neurogenesis in human cortical organoids (Figures 3I and 5B, C). Of note, ADD1 KO in organoids did not result in a specific depletion of neurons, but rather to a reduction of the entire organoid size with a strong decrease in various NPCs and an increase in the abundance of aberrant neurons identified in the “Neurons 3” cluster. The aberrant neurons express multiple markers of ER stress, suggesting that their maturation is impaired. This could be due to (1) direct effects of ADD1 KO in neurons or (2) upstream effects in NPCs which then results in aberrant production of neurons, or due to both events. The first scenario is in line with both the general role of ADDs in maintaining cell morphology and fundamental importance of ER proteostasis for neuronal migration, maturation and neuritogenesis (Martinez et al., 2018; Vasquez et al., 2022). Instead, the second scenario is supported by our data on altered proliferation and division mode of NPCs, which in turn can lead to production of immature daughter cells. Furthermore, it has been shown that unfolded protein response (UPR) regulates fate specification of NPCs and that interfering with ER stress and UPR leads to direct neurogenesis from RG as well as microcephaly (Laguesse et al., 2015). It is thus interesting to note that our single-cell fate mapping of aberrant neurons in ADD1 KO organoids, shows differentiation trajectory more likely to emerge directly from RG (Figure S9G, H), which would be an indication of potential direct neurogenesis. Together with the reduced size of ADD1 KO organoids (Figure 4C), these data suggest that aberrant neurogenesis might have origin in the biology of RG. Finally, it is possible that ADDs have effects in both progenitors and neurons and it would be interesting to examine the cell-intrinsic roles of ADDs in neuronal maturation and migration at various time points of human cortical development.

### ADDs in the evolutionary expansion of neocortex and neurodevelopmental pathologies

In mouse, OE of both ADD1 and ADD2 individually, led to an increase in the abundance of BPs, whereas OE of ADD3 did not, unless ADD1 was co-expressed as well (Figure 2 and S3). In contrast, depletion of ADD3 in human fetal cortex does lead to a decrease in BPs (Kalebic et al., 2019). This suggest that although ADD3 is not sufficient to promote BP abundance in mouse, it is required for the abundance of human BP. Hence, there might be different roles of specific ADDs in various mammals. In this context, it will be interesting to study the regulation of expression and activation of various ADDs in different model systems to explore potential differences. A first insight is given by the analyses of the mRNA and protein expression patterns (Figure S1) which shows that ADD3 is enriched in both VZ and SVZ in ferret and human, whereas only in the VZ of mouse. Such pattern is common for genes associated with NPC proliferation (Florio et al., 2015) as human and ferret BPs in the SVZ have a greater proliferative capacity than the mouse BPs (Kalebic and Huttner, 2020; Lui et al., 2011). Furthermore, only ADD3, and not other ADDs, has been strongly implicated in glioblastoma (Barelli et al., 2024; Kiang et al., 2020; Rani et al., 2013), the major human brain tumor, whose stem cells bear a striking resemblance to developmental bRG (Bhaduri et al., 2020).

Besides the role in adult tumors of the brain, ADDs have been implicated in various neurodevelopmental pathologies, such as schizophrenia, cortical malformations and cerebral palsy (Bosia et al., 2016; Kruer et al., 2013; Qi et al., 2022). ADD1 variants have been associated with microcephaly and ventriculomegaly as well as corpus callosum dysgenesis (Qi et al., 2022). Considering our findings, it is tempting to speculate that cortical malformations due to ADD1 loss of function are linked to its effects on the mitotic spindle orientation. Accordingly, mutations in proteins associated with the spindle orientation are often found in microcephaly (Bond and Woods, 2006; Gilmore and Walsh, 2013; Ossola and Kalebic, 2021; Sun and Hevner, 2014). However, these mutations typically don’t lead to strong impairment of mouse neurogenesis, but can be recapitulated only in species with an already expanded neocortex (Johnson et al., 2018), suggesting a stronger effect of the spindle orientation in such species. Interestingly, ADD1 KO mice did not recapitulate the microcephaly phenotype, although they exhibited ventriculomegaly and corpus callosum dysgenesis (Qi et al., 2022). In contrast, we show that depletion of ADDs in human cortical organoids through ADD1 KO led to a strikingly reduced organoid size (Figure 4C).

Although malformation-causing variants of ADD1 are associated with its binding to ADD2, the ADD2 KO mice do not show any of the malformation phenotypes (Gilligan et al., 1999), suggesting again that ADD1 is the obligatory partner of the ADDs complex, which is in agreement with our results. ADD3 variants have been associated with inherited cerebral palsy (Kruer et al., 2013; Sanchez Marco et al., 2022). The mutation, that likely affects its binding to ADD1, led to an impairment of the actin-capping function of adducin, which resulted in abnormal proliferation of cultured patient fibroblasts (Kruer et al., 2013). This further suggests an important role of ADD3 in human brain development and the related pathologies.

Taken together, we propose that ADDs present important morpho-regulatory proteins that underlie proliferation of BPs and fate of NPCs, which may have implications to neocortical evolution and human neurodevelopmental pathologies.

## Acknowledgements

We are grateful to the services and facilities of HT for the outstanding support provided, notably, N. Maghelli and F. Casagrande from the National facility (NF) for Light Imaging, C. Peano and the team of the NF for Genomics, M.Bonfanti and D. dalle Nogare from the NF for Data Handling and Analysis and the team of the NF for Genome engineering and Disease modeling. We are thankful to J.Helppi, Katrin Reppe and the team of the Biomedical Services of MPI-CBG Dresden for outstanding support with ferret experiments. NK is very grateful to Wieland Huttner (MPI-CBG) for his support. We thank T. Namba (HiLife) for the critical reading of the manuscript and all members of the Kalebic lab for helpful discussions. CO, NC, EC are Ph.D. students within the European School of Molecular Medicine (SEMM). Research in the Kalebic lab is supported by funds from HT and grants from AIRC (MFAG 2022 ID 27157) and Gilbert Family Foundation (#923004) to NK.

## Author contributions

Conceptualization: NK; Methodology: CO, NC, IB, CA, GF; Formal analysis: CO, NC, SF, EC; Investigation: CO, EC, IB, NK; Data curation: CO, NC; Writing – original draft: NK; Writing – review and editing: all authors; Visualization: CO, NC, SF, EC, CA, NK; Supervision: NK; Project administration: NK; Funding acquisition: NK.

## Materials and methods

| Reagent/Resource | Reference or Source | Identifier or Catalog Number |
| --- | --- | --- |
| **Experimental Models** |  |  |
| H9 cell line (human) | WiCell | WA009 |
| Mouse: C57Bl/6J | Charles River | N/A |
| Ferret *Mustela putorius furo* | Marshall Bioresources | N/A |
| **Recombinant DNA** |  |  |
| pCAG | (Kalebic et al., 2019) | N/A |
| pCAG-GFP | (Kalebic et al., 2019) | N/A |
| pCAG-LynGFP | (Kalebic et al., 2019) | N/A |
| pGEM-T Easy Vector Systems | Promega | A1360 |
| pGEM-ADD1 | This paper | N/A |
| pCAG-ADD1 | This paper | N/A |
| pCAG-ADD2 | This paper | N/A |
| pCAG-ADD3 | This paper | N/A |
| pUC19 | Addgene | #50005 |
| SpCas9 | Addgene | #48137 |
| pEGFP-puro | Addgene | #45561 |
| **Oligonucleotides** |  |  |
| ADDs cloning primers, see Table S1 | This paper | N/A |
| **Commerical kits** |  |  |
| RNeasy Mini Kit RNA purification | QIAGEN | 74104 |
| SuperScript IV Reverse Transcriptase | ThermoFisher | 18090010 |
| QIAquick PCR Purification Kit | QIAGEN | 28104 |
| EndoFree Plasmid Maxi kit | QIAGEN | 12362 |
| QIAquick gel extraction kit | QIAGEN | 28704 |
| QuickExtract DNA Extraction Solution | Epicentre | #QE09050 |
| NucleoSpin Gel and PCR clean-up | Macherey-Nagel | 740609.50 |
| Neural Tissue Dissociation Kit (P) | Miltenyi | 130-092-628 |
| Click-iT™ Plus EdU Alexa Fluor™ 647 | Invitrogen | C10340 |
| **Bacterial strains** |  |  |
| XL-10 Gold Ultracompetent cells | Agilent | 200315 |
| **Antibodies** |  |  |
| Rabbit anti-panADD1 polyclonal antibody (IF 1:200) | Abcam | Ab51130 |
| Rabbit anti-ADD1 (phospho S726) monoclonal antibody (IF 1:200) | Abcam | ab40760 |
| Mouse anti-ADD2 monoclonal antibody (IF 1:200) | Santa Cruz | sc-376063 |
| Rabbit anti-ADD3 polyclonal antibody (IF 1:200) | Atlas Antibodies | HPA035696 |
| Rabbit anti-Ki67 monoclonal antibody (IF 1:200) | Cell Signalling | 9129S |
| Goat anti-Sox2 polyclonal antibody (IF 1:200) | BioTechne | AF2018 |
| Rat anti-CTIP2 monoclonal antibody (IF 1:500) | Abcam | ab18465 |
| Mouse anti-SATB2 monoclonal antibody (IF 1:200) | Abcam | ab51502 |
| Rabbit anti-HuCHuD monoclonal antibody (IF 1:200) | Abcam | ab184267 |
| Chicken anti-GFP polyclonal antibody (IF 1:1000) | Abcam | ab13970 |
| Rabbit anti-Tbr2 monoclonal antibody (IF 1:400) | Abcam | ab183991 |
| Mouse anti-ZO1 monoclonal antibody (1:500 IF) | Invitrogen | MA3-39100-A647 |
| Mouse anti-p(ser55) Vimentin monoclonal antibody (1:500 IF) | MBL Company | D076-3 |
| Mouse anti-Actin monoclonal antibody (WB 1:20.000) | Millipore-Sigma | MAB15 |
| Donkey anti-Rabbit Alexa Fluor PLUS 488 (IF 1:500) | Thermofisher | A32790 |
| Donkey anti-Rabbit Alexa Fluor PLUS 555 (IF 1:500) | Thermofisher | A32794 |
| Donkey anti-Rabbit Alexa Fluor PLUS 647 (IF 1:500) | Thermofisher | A32795 |
| Donkey anti-Mouse Alexa Fluor PLUS 488 (IF 1:500) | Thermofisher | A32766 |
| Donkey anti-Mouse Alexa Fluor PLUS 555 (IF 1:500) | Thermofisher | A32773 |
| Donkey anti-Mouse Alexa Fluor PLUS 647 (IF 1:500) | Thermofisher | A32787 |
| Donkey anti-Rat Alexa Fluor PLUS 488 (IF 1:500) | Thermofisher | A48269 |
| Donkey anti-Rat Alexa Fluor PLUS 555 (IF 1:500) | Thermofisher | A48270 |
| Donkey anti-Rat Alexa Fluor PLUS 647 (IF 1:500) | Thermofisher | A48272 |
| Donkey anti-Chicken Alexa Fluor PLUS 488 (IF 1:500) | Jackson Immuno Research | J1703545155 |
| Donkey anti-Goat Alexa Fluor PLUS 488 (IF 1:500) | Thermofisher | A32814 |
| Donkey anti-goatAlexa Fluor PLUS 555 (IF 1:500) | Thermofisher | A32816 |
| Donkey anti-goatAlexa Fluor PLUS 647 (IF 1:500) | Thermofisher | A32849 |
| DAPI | Invitrogen | 62248 |
| **Chemicals, Enzymes and other reagents** |  |  |
| Vitronectin (VTN-N)<br>Recombinant Human Protein | ThermoFisher | A14700 |
| Dorsomorphin 5mM | StemCell<br>Technologies | 72102 - 10mg |
| A83-01 5mM | StemCell<br>Technologies | 72022 - 50mg |
| CHIR-99021 10mM | Tocris | 4423 - 10mg |
| SB-431542 10mM | StemCell<br>Technologies | 72232 - 10mg |
| Matrigel | Corning | FLC354277 |
| N2 supplement | Life Technologies | 17502-048 |
| B27 supplement | Life Technologies | 17504-044 |
| Lipofectamine | ThermoFisher | STEM008 |
| Lipod293 transfecting reagent | Signagen | SL100668 |
| Stemflex | ThermoFisher | A3349401 |
| Cas9 protein (Alt-R® S.p. HiFi<br>Cas9 Nuclease V3) | IDT | 1081061 |
| BG iMatrix-511 Silk | Tebubio | 348RL511S |
| Bolt 4-12% Bis-Tris Plus Gels | ThermoFisher | NW04120BOX |
| Trans-Blot Turbo Transfer<br>System | BioRad | 1704150 |
| **Software and Data** |  |  |
| Fiji ImageJ | v2.9.0 | https://imagej.net/software/fiji/ |
| GraphPad Prism | v9.1.2 | https://www.graphpad.com/features |
| Zeiss Zen Blue | v3.7 | https://www.zeiss.com/microscopy/en/products/software/zeiss-zen.html |
| Plotnine | v0.12.4 | https://plotnine.org/ |
| Seaborn | v0.12.2 | https://seaborn.pydata.org/ |
| Matplotlib | v3.7.3 | https://matplotlib.org/ |
| Sanbomics | v0.1.0 | https://github.com/mousepixels/sanbomics |
| Scanpy | v1.9.6 | https://scanpy.readthedocs.io/en/stable/ |
| Scrublet | v0.2.3 | https://github.com/wolock/scrublet |
| PyEnsembl | v.2.3.13 | https://github.com/openvax/pyensembl |
| Clustree | v0.2.1 | https://github.com/lazappi/clustree |
| Decoupler | v1.5.0 | https://decoupler-py.readthedocs.io/en/latest/ |
| PyDESeq2 | v0.4.1 | https://pydeseq2.readthedocs.io/en/latest/ |
| GSEAPY | v1.0.6 | https://gseapy.readthedocs.io/en/latest/introduction.html |
| scCODA | v0.1.9 | https://sccoda.readthedocs.io/en/latest/ |
| CellRank | v2.0.5 | https://cellrank.readthedocs.io/en/latest/ |

### Experimental animals: mice

All experimental procedures were conducted in agreement with the Italian Legislation after approval by the Ministry of Health (393/2021-PR). Animals used for this study arrived from Charles River and were kept in standardized hygienic conditions at Cogentech animal house in Milan, with free access to food and water, in 12h / 12h light/dark cycle. All experiments were performed on C57Bl/6J mice, in the dorsolateral telencephalon of mouse embryos, at a medial position along the rostro-caudal axis. The sex of embryos was not determined as it is not likely to be of relevance for the results obtained in the present study. Embryonic day E0.5 was set at noon of the day of vaginal plug identification.

### Experimental animals: ferrets

All experimental procedures were conducted in agreement with the German Animal Welfare Legislation after approval by the Landesdirektion Sachsen (TVV03/2020). Timed-pregnant ferrets were obtained from Marshall Bioresources (NY, USA) and housed at the Biomedical Services Facility (BMS) of the MPI-CBG in Dresden with free access to food and water, in 16h / 8h light/dark cycle. All experiments were performed in the dorsal telencephalon at a medial position along the rostro-caudal axis. The sex of embryos was not determined as it is not likely to be of relevance for the results obtained in the present study. Observed mating date was set to E0.

## METHODS DETAILS

### Plasmids

ADDs cDNA was obtained from human cortical organoids. ADD1 (sequence ENST00000398129.5) was first amplified by PCR and cloned into pGEM-Teasy Vector System (Table S1); consequently, it was subcloned into SCAG plasmid using SphI and SacI restriction enzymes. ADD2 (sequence ENST00000264436.9) and ADD3 (sequence ENST00000356080.9) were amplified by PCR with primers that introduced restriction sites at both ends (Table S1) and cloned into pCAGGS using XhoI and BglII restriction enzymes. After ligation, DNA was used to transform competent bacteria (XL-10 Gold). The correct sequence was confirmed by Sanger sequencing.

### In vivo experiments in mouse

In utero electroporations (IUE) were performed as previously described (Kalebic et al., 2016). E13.5 pregnant mice were anesthetized with isofluorane, first in a narcosis box (with 4% isoflurane) and then with a 2.5% isoflurane mask under the hood. Animals were injected subcutaneously with the analgesic (Rimadyl); the peritoneal cavity was surgically opened and the uterus was exposed. Embryos were injected intraventricularly with a solution containing 0.1% Fast Green (Sigma), 1,5 ug/ul of one of the experimental plasmids (pCAG, pCAG-ADD1, pCAG-ADD2 or pCAG-ADD3) and 1 ug/ul of the pCAG- LynGFP vector in clean PBS. All electroporations were performed with six 50-msec pulses of 28-33 V at 1 s intervals. Once IUE was performed on all embryos, the uterus was placed back in the peritoneal cavity and muscles and skin were sutured with a 4-0 suture. Animals were kept under monitoring until they woke up from anaesthesia. For the cell cycle re-entry experiments, a single pulse of EdU was injected 24h after the IUE at day E14.5 (Kalebic et al., 2019). Mice were sacrificed by cervical dislocation and embryos were harvested at day E15.5 or E18.5.

### In vivo experiments in ferret

In utero electroporation of ferret embryos as well as pre-operative and post-operative care of ferrets were performed as previously described (Kalebic et al., 2018) and as detailed in the protocol (Kalebic et al., 2020). All the ferrets that underwent *in utero* electroporation underwent another surgery four days later that followed the same pre-operative care, anesthesia and analgesia as described previously (Kalebic et al., 2018). This second surgery was performed in order to remove the electroporated embryos by Caesarian section and to carry out a subsequent complete hysterectomy. Animals were kept at the BMS of the MPI-CBG for at least two weeks after the second surgery after which they were donated for adoption. Embryonic brains were dissected and PFA-fixed for the immunofluorescence analyses.

### In vitro experiments: transfection of HeLa cells

For testing the ability of ADDs plasmids to overexpress their insert, HeLa cells were used. HeLa cells, cultured in the same medium as HEK293T cells (see below), were seeded into 24-well plates at a density of 100,000 cells/well and transfected in suspension using Lipofectamine 2000 following the manufacturer’s instructions (ThermoFisher). Cells in each well were transfected with 0,6 ug of total DNA. Each ADDs and empty vectors as controls were transfected along with LynGFP and 24 hours after transfection cells were fixed and stained.

### In vitro experiments: transfection of HEK293T cells

For ADD1 knock-out, three sgRNAs were designed and tested for efficiency in HEK293T cells by transfection (Figure S7A). Guide RNAs were cloned inside pUC19 (Addgene) using BbsI (Thermo Fisher Scientific) restriction sites as previously described (Shalem et al., 2014). HEK293T cells were cultured in Dulbecco’s modified Eagle’s medium (DMEM, Life Technologies) supplemented with 10% fetal bovine serum (FBS, Life Technologies), 1% L-glutamine and 100U/ml antibiotics (PenStrep, Life Technologies). The cells were maintained at 37°C in a 5% CO2 humidified atmosphere. For ADD1 knock-out, 10^5^ cells/well were seeded into 24-well plates (Corning) and transfected with 3 µL Lipod293 transfecting reagent per well using 750 ng of pSpCas9, 250 ng of sgRNA plasmid and 50 ng of pEGFP-puro. Cells were cultured for 3 days before DNA extraction. For editing efficiency analysis, genomic DNA was extracted using QuickExtract DNA Extraction Solution. The target region was PCR-amplified using MyTaq HS RedMix 2X (Meridian Bioscience) using the primers listed in Table S1. PCR products were purified using NucleoSpin Gel and PCR clean-up, Sanger sequenced and analysed by Synthego’s ICE v2 CRISPR Analysis software.

### In vitro experiments: generation of H9 ADD1 KO cell lines

H9 ADD1 KO cell lines were generated at National Facility for Genome Engineering, Human Technopole, Milan, Italy. H9 cells were maintained in Stemflex culturing medium and seeded on 0,25ug/cm2 BG iMatrix-511 Silk in suspension as previously described (Fiacco et al., 2024). The cells were maintained at 37°C in a 5% CO2 humidified atmosphere. H9 pluripotent stem cells at passage 36 were nucleofected on Lonza Amaxa, 4D nucleofector according to manufacturer’s instructions. Briefly, 150 pmol of sg1ADD1 were complexed with 122 pmol of Cas9 protein (Alt-R® S.p. HiFi Cas9 Nuclease V3) and incubated at room temperature for 20 minutes to form RNPs. H9 cells were detached using Accutase (Sigma-Aldrich) and 2×10^5^ cells were nucleofected with RNP, Alt-R Cas9 Electroporation enhancer (IDT) at a final concentration of 4 µM, and 2ug of CleanCap EGFP mRNA (Tebubio). Cells were nucleofected using CB-150 program and P3 primary cell solution (Lonza). After nucleofection, cells were immediately transferred to a 12-well plate (Corning) containing pre-warmed Stemflex medium supplemented with 0,25 ug/cm2 BG iMatrix-511 Silk and CEPT mixture.

Four days after nucleofection, GFP+ cells were enriched by cell sorting. Cells were cultured for 2 additional days before DNA extraction. The bulk population of cells was serially diluted to obtain single clones. 33 single-cell clones were picked and analyzed by Sanger sequencing to check for desired editing. The selected clones (1_C11, 1_E7 and 2_B12) have different indels patterns in the two alleles, excluding for allelic dropout (Figure S7). The resulting ADD1 KO cell lines were checked for genome integrity by whole-genome sequencing (WGS) data at a 1x coverage (Low Pass Sequencing, LPS), by comparing DNA from parental and edited cell lines. Obtained results excluded the presence of large aneuploidies, chromosomal abnormalities or clinically relevant deletions or duplications (data not shown). Cell line authentication was performed by STR analysis on a Spectrum Compact CE System (Promega) using GenePrint® 10 System (Promega) (data not shown).

### In vitro experiments: H9 cell culture

Each of the three ADD1 KO cell lines were used to derive 3 independent batches of organoids along with the WT H9 as a control. H9 ESCs were cultured at 37 °C and 5% CO2 in TeSR-E8 medium in 10 cm Petri dish coated with Vitronectin. The medium was changed daily and cells were split using 0.5 mM EDTA when 80% confluent. They were frozen in StemCell Banker medium and thawed in fresh TeSR-E8 + Rock Inhibitor.

### In vitro experiments: generation of cortical organoids

Cortical organoids were derived according to the sliced organoid protocol (Qian et al., 2020). Embryoid bodies (EBs) were generated from H9 colonies incubated in the neural induction medium (Medium 1) consisting of 20% Knockout serum replacement (Gibco), 1X GlutaMAX (Gibco), 1X MEM-NEAA (Gibco), 1X Pen/Strep (Gibco), 2 µM Dorsomorphin (Stem cell technologies), 2 µM A83-01 (Stem cell technologies), 1X 2-Mercaptoethanol (Gibco) in DMEM:F12 (Gibco). Medium was changed on days 1, 3 and 4. From day 7 to day 14, EBs were embedded in Matrigel (Corning) and patterned towards a forebrain fate in Medium 2 consisting of DMEM:F12 (Gibco), 1X N2 Supplement (Life Technologies), 1X Pen/Strep, 1X MEM-NEAA, 1X GlutaMax, 1 μM CHIR-99021 (Tocris), and 1 μM SB-431542 (Stem cell technologies). On day 14, embedded organoids were mechanically removed from Matrigel and incubated in the differentiation medium (Medium 3.1) composed of DMEM:F12, 1X N2 and 1X B27 Supplements (Life Technologies), 1X Pen/Strep, 1X 2- Mercaptoethanol, 1X MEM-NEAA, 2.5 μg/ml Human Insulin (Sigma-Aldrich) until day 35. From this time onwards organoids were left in orbital shaking at 120rpm. From day 35 to day 70, Medium 3.1 was supplemented with 1% Matrigel (Medium 3.2). At day 45, organoids were either fixed for analysis or sliced for maintenance. For slicing, organoids were embedded in 3% low melting point agarose (Invitrogen) dissolved in DMEM:F12 and sliced into 500 µm sections at 0.1 mm/s speed and 1 mm amplitude using a VT 1200S vibratome (Leica Microsystems). Medium was changed every 2 days until day 75, when the organoids were harvested for histological analysis and scRNAseq.

### Preparation of tissue and organoids for microscopy

Embryonic brains were fixed in 4% paraformaldehyde for 24h (mice) or 48h (ferret) at 4°C, dehydrated in 20% and 30% sucrose and embedded in frozen OCT. Fixed tissue was cut at a cryostat to 30 µm sections. Organoids day 45 and day 75 were fixed in 4% paraformaldehyde for 30min at RT, dehydrated in 20% and 30% sucrose and embedded in frozen OCT. Fixed tissue was cut at the cryostat to 15 µm sections.

### Immunofluorescence

Immunofluorescence was performed as previously described (Kalebic et al., 2019), including antigen retrieval, permeabilization with Triton X-100, quenching in glycine and blocking with donkey serum solution. Primary antibodies were incubated in blocking buffer over night at 4°C, secondary antibodies (1:500) and DAPI (Sigma) in blocking solution for 1-2 h at room temperature before being mounted on microscopy slides with Mowiol.

### Image acquisition and analysis

The images were acquired with a Zeiss LSM 980 or Zeiss LSM980-NLO point-scanning confocal system. PlanApo 10X/0.45 dry, PlanApo 20X/0.8 dry or PlanApo 40X/1.4NA oil immersion objectives and 405 nm, 488 nm, 561 nm, 639 nm laser lines were used. 1 µm optical sections were acquired for all experiments, except for morphological analysis, for which 0.3 µm sections were acquired. Zen Blue 3.7 (Zeiss) was used to perform tiles stitching. The parameters of acquisition were established in control conditions and kept constants for all the samples within the same experiment. We blindly analyzed each experiment using Fiji, plotted the results and performed statistical analyses with Prism (GraphPad Software). Cell quantifications were performed manually using the CellCounter function, except for the cell proliferation in d45 cortical organoids, which was assessed by automatically segmenting Sox2+, Ki67+ and Edu+ progenitors with Stardist 2D (Fiji, Figure S8D-F).

ADDs level of expression was evaluated in ferret upon ADDs KO by measuring the signal intensity on IF staining. GFP signal in control and KO ferret was segmented using Labkit (Fiji); the generated mask was overlapped on ADDs staining and the ADDs signal was quantified in areas corresponding to GFP+ cells by measuring average intensity.

Morphological analysis (Figures 3 and S5) was performed as established earlier (Kalebic et al., 2019), using the TrackEM2 Fiji plugin and Neuron Process Analysis macro. Individual GFP+Ki67+ BPs in the mouse SVZ were isolated by cropping an area of 53.3 µm x 53.3 µm surrounding the cell. Cell morphology was manually annotated on Lyn-GFP signal through Z-stack using TrackEM2 Fiji plugin. The obtained masks were exported and automatically analyzed as Z-stack max-projections using Neuron Process Analysis macro (Kalebic et al., 2019). When needed the cell body area was corrected manually. The parameters automatically quantified were the number of primary proceses (arising from the cell body) and total processes (counting of all the process extremites), the branching index (ratio between the number of all process extremities to the primary processes) and the cell body (soma) radius. In addition, a Sholl analysis was performed by counting the number of processes intersecting concentric radiuses (2r, 3r, 4r) centered on the middle of the cell body.

Spindle orientations in dividing APs (Figure 5H, I) were determined by calculating the angle between the equatorial plate and the ventricular surface (identified by ZO1) using Fiji. Posphovimentin staining was used to confirm the mitoses (data not shown). Mitotic cells adjacent to the ventricle were identified and 253 cells from 3 different d45 cortical organoid batches were analyzed. The percentage of vertical (60°-90°), oblique (30°-60°) and horizontal (0°-30°) cell divisions was calculated and compared among WT and ADD1 KO.

### Immunoblotting

WT and ADD1KO organoids (d75) were homogenized 10% (w/v) in Tris-HCl pH 8.0 50 mM, 150 mM NaCl and 1% SDS. Total proteins (20 µg) were separated on NuPAGE 4–12% gels and blotted onto nitrocellulose membranes (Trans-Blot Turbo Transfer Pack kit). After blocking with 5% dry milk for 1 h at room temperature, membranes were incubated overnight at 4°C with anti-pan Adducin1 (1:4000, AbCam) or Anti Adducin2 (1:1000, Santa Cruz) and Anti Adducin3 (1:1000, Atlas prestige), then with HRP-conjugated secondary antibodies (BioRad) 1 h at room temperature. Peroxidase activity was detected using ECL (BioRad) and visualized with a ChemiDoc imaging system (BioRad).

### Single cell RNA sequencing

Day 75 organoids from 3 batches of ADD1KO (1E7 clone) and 4 batches of WT were harvested and dissociated to perform single cell RNA sequencing analysis. 10 to 14 organoids were dissociated with the Neural Tissue Dissociation Kit (P), using gentleMACS dissociator. Total RNA was extracted from 10,000 single cells for each sample. RNA extraction and sequencing were performed by the National facility for Genomics (Human Technopole). Single cells were encapsulated into Gel Emulsion Droplets using the Chromium Controller system (10x Genomics) targeting 10000 cells for each sample and 3’ Gene Expression libraries were generated in accordance with the 10x Genomics Protocols (Chromium Single Cell 3’ Reagent Kits, v3.1 Chemistry Dual Index). Libraries were pooled at a ratio dependent on the 10x Genomics indication and loaded on the Novaseq6000 sequencer to achieve a minimum of 50’000 reads/cell for each library. All downstream analyses have been performed with Scanpy single cell framework. Raw count matrices were imported from CellRanger and subjected to pre-processing and analytical steps described below.

### Transcriptomic analysis

Data pre-processing included a total of 119,666 cells. Neotypic doublets were identified and removed for each sample independently using the scrublet package. Next, samples were concatenated and subjected to quality control to remove low quality cells and insufficiently abundant genes. Cells were filtered based on counts, number of expressed genes and the fraction of mitochondrial and ribosomal genes using scanpy.pp.filter_cells and scanpy.pp.filter_genes as follows: min_counts > 5500; max_counts < 30000; min_genes > 2500; max_genes < 8500; % mito genes < 10; % ribo genes < 30. In addition, only genes present in at least 50 cells were kept for downstream processing. PyEnsembl, python interface to Human Reference Genome (GRCh38) was used to identify and keep protein coding genes, while removing non-coding genes. After the pre-processing, dataset contained 69,671 high quality cells and 17,001 protein-coding genes.

Raw counts were normalized, log-transformed and scaled using the scanpy.pp.normalize_total, sc.pp.log1p and sc.pp.scale, respectively. Highly variable genes (HVGs) were identified in a batch-wise manner (batch_key = sequencing round) using sc.pp.highly_variable_genes, following default set of parameters (min_disp=0.5, min_mean=0.0125 and max_mean=3).

Principal component analysis (PCA) was computed based on the HVGs from the previous step, using the scanpy.tl.pca, and top 50 PCs were initially retained. Integration of individual samples was performed using the Scanpy’s implementation of Harmony, with the information of the sequencing round as a batch covariate. The nearest neighbors graph was computed using the first 30 PCs and range of numbers of nearest neighbors (NN = 20, 30, 40, 50, 80, 100) to identify the optimal balance in local and global structure preservation. Eventually, NN = 80 was used for downstream steps.

Clustering was performed using the Scanpy implementation of the Leiden algorithm (scanpy.tl.leiden). Initially, data was clustered at low resolution (R = 0.1) to identify major cell populations. This approach identified biologically expected cell clusters (radial glia, intermediate progenitors and neurons) and two clusters of od-target cell populations, which showed increased gene expression of non-telencephalic marker genes. Hence, cell cluster scoring for the average expression of the selected marker genes (LHX9, MSX1, NHLH2, SLC17A6, GBX2, SHOX2, RSPO3 and OTX2) was performed using the scanpy.tl.score_genes, with the reference set of genes (n = 50) randomly sampled from the dataset. This resulted in od-target clusters scoring significantly higher for selected marker genes compared to the rest of the clusters and they were therefore removed from the downstream analyses. Following the re-normalization and re-computing of HVGs and PCs, data was re-integrated (Harmony) and nearest neighbor graph was re-calculated (n_pcs = 30, n_neighbors = 40). Lastly, clustering was repeated (R = 0.18) to obtain the final cell clusters. Optimal clustering resolution was determined using the clustree package in R. Clusters were annotated by analyzing the expression of known marker genes (see Figure S9A for the list of markers).

Pseudo-bulk replicates were generated by summarizing raw gene counts across cells per sample and per cell cluster using the decoupler.get_pseudobulk function from the decoupler framework. Pseudo-bulk replicates with less than 10 cells and 10k counts were discarded prior to the downstream analysis. Raw pseudo-bulk counts were normalized and log-transformed and HVGs and PCs were computed as before. Insufficiently abundant genes were removed using the decoupler.plot_filter_by_expr function (min_count = 5, min_total_count =10, min_prop = 0.7) for all the clusters individually. Pseudo-bulked individual clusters were analyzed for differentially expressed genes (DEGs) using the PyDESeq2 package. Genes with log2FC > 0.58 (corresponding to FC > 1.5) and p_adjusted < 0.05 (correction method: Benjamini-Hochberg) were considered differentially expressed, except for the comparison of Neurons 3 vs Neurons 2, in which the genes were considered differentially expressed if p_adjusted < 0.05 and log2FC > 1.

Gene ontology (GO) analysis was performed using the GSEAPY package separately for up-regulated and down-regulated genes for each cell cluster. GO lists were downloaded from the Molecular Signature Database website (GSEA | MSigDB (gsea-msigdb.org). Background gene lists were comprised of the genes used for the DeSeq2 modeling. Gene ontology terms were considered significantly enriched if p_adjusted < 0.05. Compositional analysis was performed using the scCODA framework. Samples with the atypical distribution of cell clusters were not included in the analysis. Information on the experimental condition (WT vs KO) was the only covariate used for the modeling.

Pseudotime analysis was performed using the scanpy.tl.dpt function, following the default set of parameters. The cell with the highest expression of TOP2A was selected as a root cell. Generalized Perron Cluster Cluster Analysis (GPCCA) was used to compute the macrostates (GPCCA.predict_terminal_states) and fate probabilities towards those macrostates (GPCCA.compute_fate_probabilities), as part of the CellRank framework, identifying thus the cycling progenitors and Neurons 1 terminal states. Instead, Neurons 3 terminal state was recovered in an analogous way, by selecting 30 cells with highest expression of PERK, a key marker involved in ER stress. Aggregated fate probabilities per progenitor clusters (either IP or RG) were then computed as previously described (Lange et al., 2022).

### Statistical analysis

For the results shown in Figures 1-4, 5H, I, S1-8 the statistics analyses were conducted on Prism. For the transcriptomic data the statistics analyses were conducted in scanpy-based frameworks. To test for statistical significance, we used Welch’s *t* test, Student’s *t* test, Kolmogorov-Smirnov test and two-way ANOVA. For each graph, the number of samples, statistical test and the p value are noted in the respective figure legends.

## References

Arai, Y., Pulvers, J.N., Haffner, C., Schilling, B., Nusslein, I., Calegari, F., and Huttner, W.B. (2011). Neural stem and progenitor cells shorten S-phase on commitment to neuron production. Nat Commun 2, 154.

Barelli, C., Kaluthantrige Don, F., Iannuzzi, R.M., Bertani, I., Osei, I., Sorrentino, S., Villa, G., Sokolova, V., Iorio, F., and Kalebic, N. (2024). Morphoregulatory ADD3 underlies glioblastoma growth and forma9on of tumor-tumor connections. bioRxiv.

Barkalow, K.L., Italiano, J.E., Jr., Chou, D.E., Matsuoka, Y., Bennett, V., and Hartwig, J.H. (2003). Alpha-adducin dissociates from F-actin and spectrin during platelet activation. J Cell Biol 161, 557–570.

Bennett, V., Gardner, K., and Steiner, J.P. (1988). Brain adducin: a protein kinase C substrate that may mediate site-directed assembly at the spectrin-actin junction. The Journal of biological chemistry 263, 5860–5869.

Bhaduri, A., Di Lullo, E., Jung, D., Muller, S., Crouch, E.E., Espinosa, C.S., Ozawa, T., Alvarado, B., Spatazza, J., Cadwell, C.R., et al. (2020). Outer Radial Glia-like Cancer Stem Cells Contribute to Heterogeneity of Glioblastoma. Cell Stem Cell 26, 48–63 e46.

Bond, J., and Woods, C.G. (2006). Cytoskeletal genes regulating brain size. Curr Opin Cell Biol 18, 95–101.

Bosia, M., Pigoni, A., Zagato, L., Merlino, L., Casamassima, N., Lorenzi, C., Pirovano, A., Smeraldi, E., Manunta, P., and Cavallaro, R. (2016). ADDing a piece to the puzzle of cognition in schizophrenia. Eur J Med Genet 59, 26–31.

Chan, P.C., Hsu, R.Y., Liu, C.W., Lai, C.C., and Chen, H.C. (2014). Adducin-1 is essential for mitotic spindle assembly through its interaction with myosin-X. J Cell Biol 204, 19–28.

Chen, C.L., Hsieh, Y.T., and Chen, H.C. (2007). Phosphorylation of adducin by protein kinase Cdelta promotes cell motility. J Cell Sci 120, 1157–1167.

Cubillos, P., Ditzer, N., Kolodziejczyk, A., Schwenk, G., Hoffmann, J., Schutze, T.M., Derihaci, R.P., Birdir, C., Kollner, J.E., Petzold, A., et al. (2024). The growth factor EPIREGULIN promotes basal progenitor cell proliferation in the developing neocortex. The EMBO journal 43, 1388–1419.

Del-Valle-Anton, L., and Borrell, V. (2022). Folding brains: from development to disease modeling. Physiol Rev 102, 511–550.

di Pietro, F., Echard, A., and Morin, X. (2016). Regulation of mitotic spindle orientation: an integrated view. EMBO Rep 17, 1106–1130.

Dong, L., Chapline, C., Mousseau, B., Fowler, L., Ramsay, K., Stevens, J.L., and Jaken, S. (1995). 35H, a sequence isolated as a protein kinase C binding protein, is a novel member of the adducin family. The Journal of biological chemistry 270, 25534–25540.

Ferent, J., Zaidi, D., and Francis, F. (2020). Extracellular Control of Radial Glia Proliferation and Scaffolding During Cortical Development and Pathology. Front Cell Dev Biol 8, 578341.

Fiacco, E., Landi, S., Zasso, J., Ambrosini, C., and Faga, G. (2024). Optimized and Scalable Precoating-Free Reprogramming of Human Peripheral Blood Mononuclear Cells into iPSCs. Curr Protoc 4, e979.

Fietz, S.A., Lachmann, R., Brandl, H., Kircher, M., Samusik, N., Schroder, R., Lakshmanaperumal, N., Henry, I., Vogt, J., Riehn, A., et al. (2012). Transcriptomes of germinal zones of human and mouse fetal neocortex suggest a role of extracellular matrix in progenitor self-renewal. Proceedings of the National Academy of Sciences of the United States of America 109, 11836–11841.

Fish, J.L., Kennedy, H., Dehay, C., and Huttner, W.B. (2008). Making bigger brains - the evolution of neural-progenitor-cell division. J Cell Sci 121, 2783–2793.

Florio, M., Albert, M., Taverna, E., Namba, T., Brandl, H., Lewitus, E., Haffner, C., Sykes, A., Wong, F.K., Peters, J., et al. (2015). Human-specific gene ARHGAP11B promotes basal progenitor amplification and neocortex expansion. Science 347, 1465–1470.

Gardner, K., and Bennett, V. (1987). Modulation of spectrin-actin assembly by erythrocyte adducin. Nature 328, 359–362.

Gilardi, C., and Kalebic, N. (2021). The Ferret as a Model System for Neocortex Development and Evolution. Front Cell Dev Biol 9, 661759.

Gilligan, D.M., Lozovatsky, L., Gwynn, B., Brugnara, C., Mohandas, N., and Peters, L.L. (1999). Targeted disruption of the beta adducin gene (Add2) causes red blood cell spherocytosis in mice. Proc Natl Acad Sci U S A 96, 10717–10722.

Gilmore, E.C., and Walsh, C.A. (2013). Genetic causes of microcephaly and lessons for neuronal development. Wiley Interdiscip Rev Dev Biol 2, 461–478.

Gonzalez-Fernandez, E., Fan, L., Wang, S., Liu, Y., Gao, W., Thomas, K.N., Fan, F., and Roman, R.J. (2022). The adducin saga: pleiotropic genomic targets for precision medicine in human hypertension-vascular, renal, and cognitive diseases. Physiol Genomics 54, 58–70.

Hsu, W.H., Wang, W.J., Lin, W.Y., Huang, Y.M., Lai, C.C., Liao, J.C., and Chen, H.C. (2018). Adducin-1 is essential for spindle pole integrity through its interaction with TPX2. EMBO Rep 19.

Johnson, M.B., Sun, X., Kodani, A., Borges-Monroy, R., Girskis, K.M., Ryu, S.C., Wang, P.P., Patel, K., Gonzalez, D.M., Woo, Y.M., et al. (2018). Aspm knockout ferret reveals an evolutionary mechanism governing cerebral cortical size. Nature 556, 370–375.

Joshi, R., Gilligan, D.M., Otto, E., McLaughlin, T., and Bennett, V. (1991). Primary structure and domain organization of human alpha and beta adducin. J Cell Biol 115, 665–675.

Juric-Sekhar, G., and Hevner, R.F. (2019). Malformations of Cerebral Cortex Development: Molecules and Mechanisms. Annu Rev Pathol 14, 293–318.

Kalebic, N., Gilardi, C., Albert, M., Namba, T., Long, K.R., Kostic, M., Langen, B., and Huttner, W.B. (2018). Human-specific ARHGAP11B induces hallmarks of neocortical expansion in developing ferret neocortex. eLife 7: e41241.

Kalebic, N., Gilardi, C., Stepien, B., Wilsch-Brauninger, M., Long, K.R., Namba, T., Florio, M., Langen, B., Lombardot, B., Shevchenko, A., et al. (2019). Neocortical expansion due to increased proliferation of basal progenitors is linked to changes in their morphology. Cell Stem Cell 24, 535–550.

Kalebic, N., and Huttner, W.B. (2020). Basal Progenitor Morphology and Neocortex Evolution. Trends Neurosci.

Kalebic, N., Langen, B., Helppi, J., Kawasaki, H., and Huttner, W.B. (2020). In Vivo Targeting of Neural Progenitor Cells in Ferret Neocortex by In Utero Electroporation. J Vis Exp.

Kalebic, N., Long, K., and Huttner, W.B. (2017). Neocortex expansion in development and evolution: the cell biology of neural stem and progenitor cells and the impact of human-specific gene expression. In Evolution of nervous systems 2e, J. Kaas, ed. (Oxford: Elsevier), pp. 73–89.

Kalebic, N., and Namba, T. (2021). Inheritance and flexibility of cell polarity: a clue for understanding human brain development and evolution. Development 148.

Kalebic, N., Taverna, E., Tavano, S., Wong, F.K., Suchold, D., Winkler, S., Huttner, W.B., and Sarov, M. (2016). CRISPR/Cas9-induced disruption of gene expression in mouse embryonic brain and single neural stem cells in vivo. EMBO reports 17, 338–348.

Kiang, K.M., and Leung, G.K. (2018). A review on adducin from functional to pathological mechanisms: future direction in cancer. Biomed Res Int 2018, 3465929.

Kiang, K.M., Zhang, P., Li, N., Zhu, Z., Jin, L., and Leung, G.K. (2020). Loss of cytoskeleton protein ADD3 promotes tumor growth and angiogenesis in glioblastoma multiforme. Cancer Lett 474, 118–126.

Kruer, M.C., Jepperson, T., Dutta, S., Steiner, R.D., Cottenie, E., Sanford, L., Merkens, M., Russman, B.S., Blasco, P.A., Fan, G., et al. (2013). Mutations in gamma adducin are associated with inherited cerebral palsy. Ann Neurol 74, 805–814.

Kuhlman, P.A., Hughes, C.A., Bennett, V., and Fowler, V.M. (1996). A new function for adducin. Calcium/calmodulin-regulated capping of the barbed ends of actin filaments. The Journal of biological chemistry 271, 7986–7991.

Kunda, P., and Baum, B. (2009). The actin cytoskeleton in spindle assembly and positioning. Trends Cell Biol 19, 174–179.

Laguesse, S., Creppe, C., Nedialkova, D.D., Prevot, P.P., Borgs, L., Huysseune, S., Franco, B., Duysens, G., Krusy, N., Lee, G., et al. (2015). A Dynamic Unfolded Protein Response Contributes to the Control of Cortical Neurogenesis. Dev Cell 35, 553–567.

LaMonica, B.E., Lui, J.H., Hansen, D.V., and Kriegstein, A.R. (2013). Mitotic spindle orientation predicts outer radial glial cell generation in human neocortex. Nat Commun 4, 1665.

Lancaster, M.A., and Knoblich, J.A. (2012). Spindle orientation in mammalian cerebral cortical development. Curr Opin Neurobiol 22, 737–746.

Lange, M., Bergen, V., Klein, M., Setty, M., Reuter, B., Bakhti, M., Lickert, H., Ansari, M., Schniering, J., Schiller, H.B., et al. (2022). CellRank for directed single-cell fate mapping. Nat Methods 19, 159–170.

Li, X., Matsuoka, Y., and Bennett, V. (1998). Adducin preferentially recruits spectrin to the fast growing ends of actin filaments in a complex requiring the MARCKS-related domain and a newly defined oligomerization domain. The Journal of biological chemistry 273, 19329–19338.

Lou, H., Park, J.J., Phillips, A., and Loh, Y.P. (2013). gamma-Adducin promotes process outgrowth and secretory protein exit from the Golgi apparatus. J Mol Neurosci 49, 1–10.

Lui, J.H., Hansen, D.V., and Kriegstein, A.R. (2011). Development and evolution of the human neocortex. Cell 146, 18–36.

Martinez, G., Khatiwada, S., Costa-Mattioli, M., and Hetz, C. (2018). ER Proteostasis Control of Neuronal Physiology and Synaptic Function. Trends Neurosci 41, 610–624.

Martinez-Martinez, M.A., De Juan Romero, C., Fernandez, V., Cardenas, A., Götz, M., and Borrell, V. (2016). A restricted period for formation of outer subventricular zone defined by Cdh1 and Trnp1 levels. Nat Commun 7, 11812.

Matsuoka, Y., Li, X., and Bennett, V. (1998). Adducin is an in vivo substrate for protein kinase C: phosphorylation in the MARCKS-related domain inhibits activity in promoting spectrin-actin complexes and occurs in many cells, including dendritic spines of neurons. J Cell Biol 142, 485–497.

Matsuoka, Y., Li, X., and Bennett, V. (2000). Adducin: structure, function and regulation. Cell Mol Life Sci 57, 884–895.

Matsuzaki, F., and Shitamukai, A. (2015). Cell Division Modes and Cleavage Planes of Neural Progenitors during Mammalian Cortical Development. Cold Spring Harb Perspect Biol 7, a015719.

Molnar, Z., Clowry, G.J., Sestan, N., Alzu’bi, A., Bakken, T., Hevner, R.F., Huppi, P.S., Kostovic, I., Rakic, P., Anton, E.S., et al. (2019). New insights into the development of the human cerebral cortex. J Anat 235, 432–451.

Namba, T., and Huttner, W.B. (2017). Neural progenitor cells and their role in the development and evolutionary expansion of the neocortex. Wiley Interdiscip Rev Dev Biol 6.

Ossola, C., and Kalebic, N. (2021). Roots of the Malformations of Cortical Development in the Cell Biology of Neural Progenitor Cells. Front Neurosci 15, 817218.

Penisson, M., Ladewig, J., Belvindrah, R., and Francis, F. (2019). Genes and Mechanisms Involved in the Generation and Amplification of Basal Radial Glial Cells. Frontiers in cellular neuroscience 13, 381.

Porro, F., Rosato-Siri, M., Leone, E., Costessi, L., Iaconcig, A., Tongiorgi, E., and Muro, A.F. (2010). beta-adducin (Add2) KO mice show synaptic plasticity, motor coordination and behavioral deficits accompanied by changes in the expression and phosphorylation levels of the alpha- and gamma-adducin subunits. Genes Brain Behav 9, 84–96.

Qi, C., Feng, I., Costa, A.R., Pinto-Costa, R., Neil, J.E., Caluseriu, O., Li, D., Ganetzky, R.D., Brasch-Andersen, C., Fagerberg, C., et al. (2022). Variants in ADD1 cause intellectual disability, corpus callosum dysgenesis, and ventriculomegaly in humans. Genet Med 24, 319–331.

Qian, X., Su, Y., Adam, C.D., Deutschmann, A.U., Pather, S.R., Goldberg, E.M., Su, K., Li, S., Lu, L., Jacob, F., et al. (2020). Sliced Human Cortical Organoids for Modeling Distinct Cortical Layer Formation. Cell Stem Cell 26, 766–781 e769.

Rakic, P. (2009). Evolution of the neocortex: a perspective from developmental biology. Nat Rev Neurosci 10, 724–735.

Rani, S.B., Rathod, S.S., Karthik, S., Kaur, N., Muzumdar, D., and Shiras, A.S. (2013). MiR-145 functions as a tumor-suppressive RNA by targeting Sox9 and adducin 3 in human glioma cells. Neuro Oncol 15, 1302–1316.

Robledo, R.F., Ciciotte, S.L., Gwynn, B., Sahr, K.E., Gilligan, D.M., Mohandas, N., and Peters, L.L. (2008). Targeted deletion of alpha-adducin results in absent beta- and gamma-adducin, compensated hemolytic anemia, and lethal hydrocephalus in mice. Blood 112, 4298–4307.

Rosenblatt, J., Cramer, L.P., Baum, B., and McGee, K.M. (2004). Myosin II-dependent cortical movement is required for centrosome separation and positioning during mitotic spindle assembly. Cell 117, 361–372.

Sanchez Marco, S.B., Buhl, E., Firth, R., Zhu, B., Gainsborough, M., Beleza-Meireles, A., Moore, S., Caswell, R., Stals, K., Ellard, S., et al. (2022). Hereditary spastic paraparesis presenting as cerebral palsy due to ADD3 variant with mechanistic insight provided by a Drosophila gamma-adducin model. Clin Genet 102, 494–502.

Shalem, O., Sanjana, N.E., Hartenian, E., Shi, X., Scott, D.A., Mikkelsen, T.S., Heckl, D., Ebert, B.L., Root, D.E., Doench, J.G., et al. (2014). Genome-Scale CRISPR-Cas9 Knockout Screening in Human Cells. Science 343, 84–87.

Shitamukai, A., Konno, D., and Matsuzaki, F. (2011). Oblique radial glial divisions in the developing mouse neocortex induce self-renewing progenitors outside the germinal zone that resemble primate outer subventricular zone progenitors. J Neurosci 31, 3683–3695.

Sousa, A.M.M., Meyer, K.A., Santpere, G., Gulden, F.O., and Sestan, N. (2017). Evolution of the human nervous system function, structure, and development. Cell 170, 226–247.

Sun, T., and Hevner, R.F. (2014). Growth and folding of the mammalian cerebral cortex: from molecules to malformations. Nat Rev Neurosci 15, 217–232.

Taverna, E., Götz, M., and Huttner, W.B. (2014). The cell biology of neurogenesis: toward an understanding of the development and evolution of the neocortex. Annu Rev Cell Dev Biol 30, 465–502.

Taylor, K.A., and Taylor, D.W. (1994). Formation of two-dimensional complexes of F-actin and crosslinking proteins on lipid monolayers: demonstration of unipolar alpha-actinin-F-actin crosslinking. Biophys J 67, 1976–1983.

Uzquiano, A., Gladwyn-Ng, I., Nguyen, L., Reiner, O., Gotz, M., Matsuzaki, F., and Francis, F. (2018). Cortical progenitor biology: key features mediating proliferation versus differentiation. J Neurochem 146, 500–525.

Vasquez, G.E., Medinas, D.B., Urra, H., and Hetz, C. (2022). Emerging roles of endoplasmic reticulum proteostasis in brain development. Cells Dev 170, 203781.

Woolner, S., O’Brien, L.L., Wiese, C., and Bement, W.M. (2008). Myosin-10 and actin filaments are essential for mitotic spindle function. J Cell Biol 182, 77–88.

Wynshaw-Boris, A. (2013). Spindle orientation: timing is everything. Neuron 79, 211–213.

Yaguchi, H., Yabe, I., Takahashi, H., Watanabe, M., Nomura, T., Kano, T., Matsumoto, M., Nakayama, K.I., Watanabe, M., and Hatakeyama, S. (2017). Sez6l2 regulates phosphorylation of ADD and neuritogenesis. Biochem Biophys Res Commun 494, 234–241.

