## Supplemental figures and legends for "Adducins regulate morphology and fate of neural progenitors during neocortical neurogenesis"

**Ossola et al.**

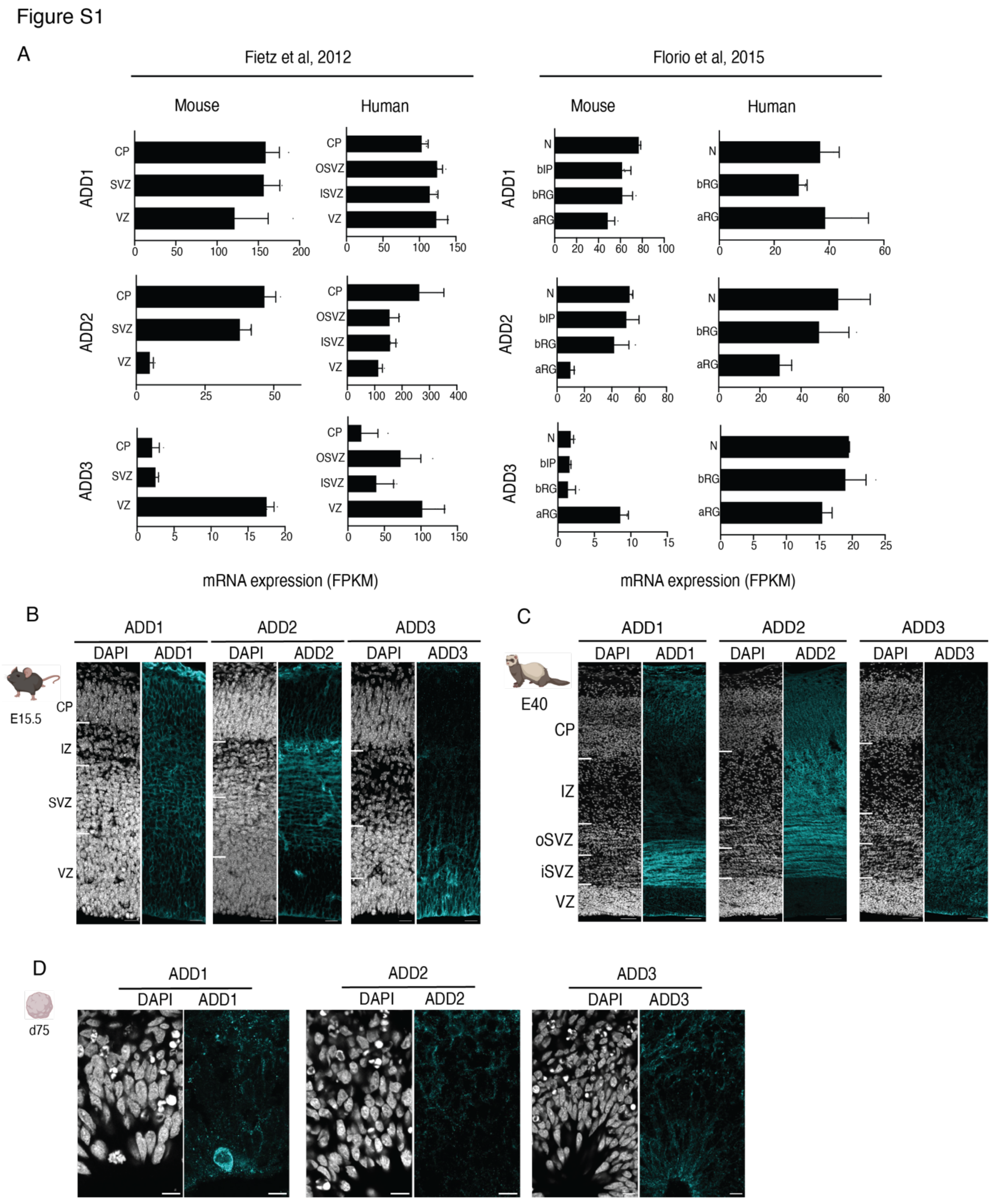


**Figure S1. Expression pattern of ADDs in mouse, ferret and human.**

(A) RNA expression pattern of ADDs according to transcriptomic data from mouse and human developing neocortex (Fietz et al., 2012; Florio et al., 2015). RNA seq data arranged for germinal zones (left panels) are from (Fietz et al., 2012). RNA seq data arranged for cell types (right panels) are from (Florio et al., 2015). ADD1 is expressed across the entire developing neocortex, from germinal zones to CP in both species. ADD2 is mainly expressed by neurons and BPs. ADD3 is enriched in NPCs. Note a higher level of expression of ADD2 and ADD3 with respect to ADD1 in human compared to mouse.

(B-D) Protein expression pattern of ADDs in mouse E15.5 neocortex (B), ferret E40 neocortex (C) and human day 75 cortical organoids differentiated according to the (Qian et al., 2020) protocol. ADD1 is expressed across the entire cortex with strong enrichment in mitotic cells (see D). ADD2 is mainly expressed in neuronal fibers and neurons. ADD3 is enriched in NPCs in the germinal zones. Scale bars 20 μm (B), 50 μm (C), 10 μm (D).

Figure S2

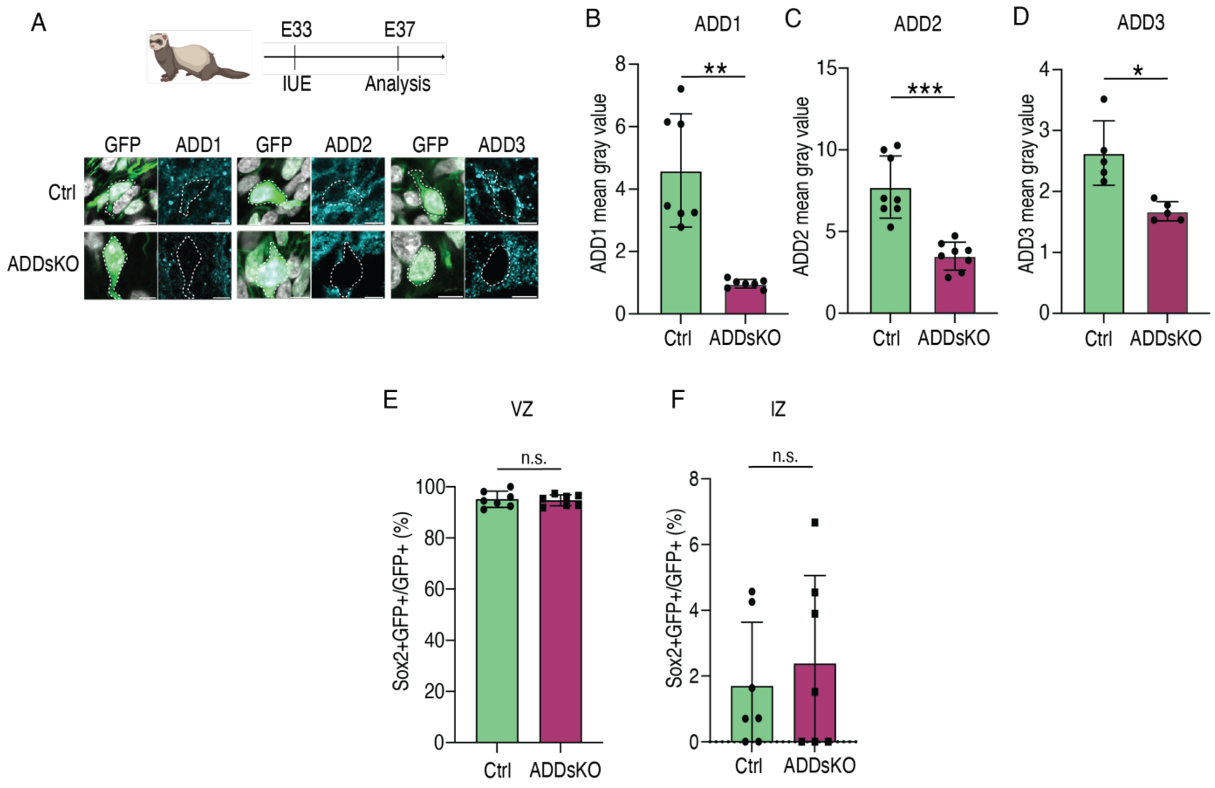


**Figure S2. Triple KO of ADDs KO in ferret developing neocortex.**

ADDs KO guides and Cas9 protein were electroporated at E33 along with GFP overexpressing plasmid; embryos were sacrificed at E37.

(A) ADDs triple KO decreases ADDs expression in GFP+ cells. DAPI staining and IF for GFP and ADDs in E37 ferret neocortex for control (upper) and ADDs KO (lower). Dashed line, examples of GFP+ cells used for the quantifications (B-D). Scale bar, 10 μm.

(B-D) ADDs triple KO effectively reduces the level of expression of ADD1 (B), ADD2 (C) and ADD3 (D) proteins. Mean intensity value measured on IF images of multiple GFP+ cells per sample. N=5-8 control and 5-8 KO embryos from 3 different litters. ***$p < 0.001$, **$p < 0.01$, *$p < 0.05$; Welch's t test.

(E-F) Triple KO of ADDs in ferret developing neocortex does not affect Sox2+ NPC abundance in VZ (E) and IZ (F). N=7 control and 7 KO embryos from 3 different litters. n.s. not statistically significant, Student's t test.

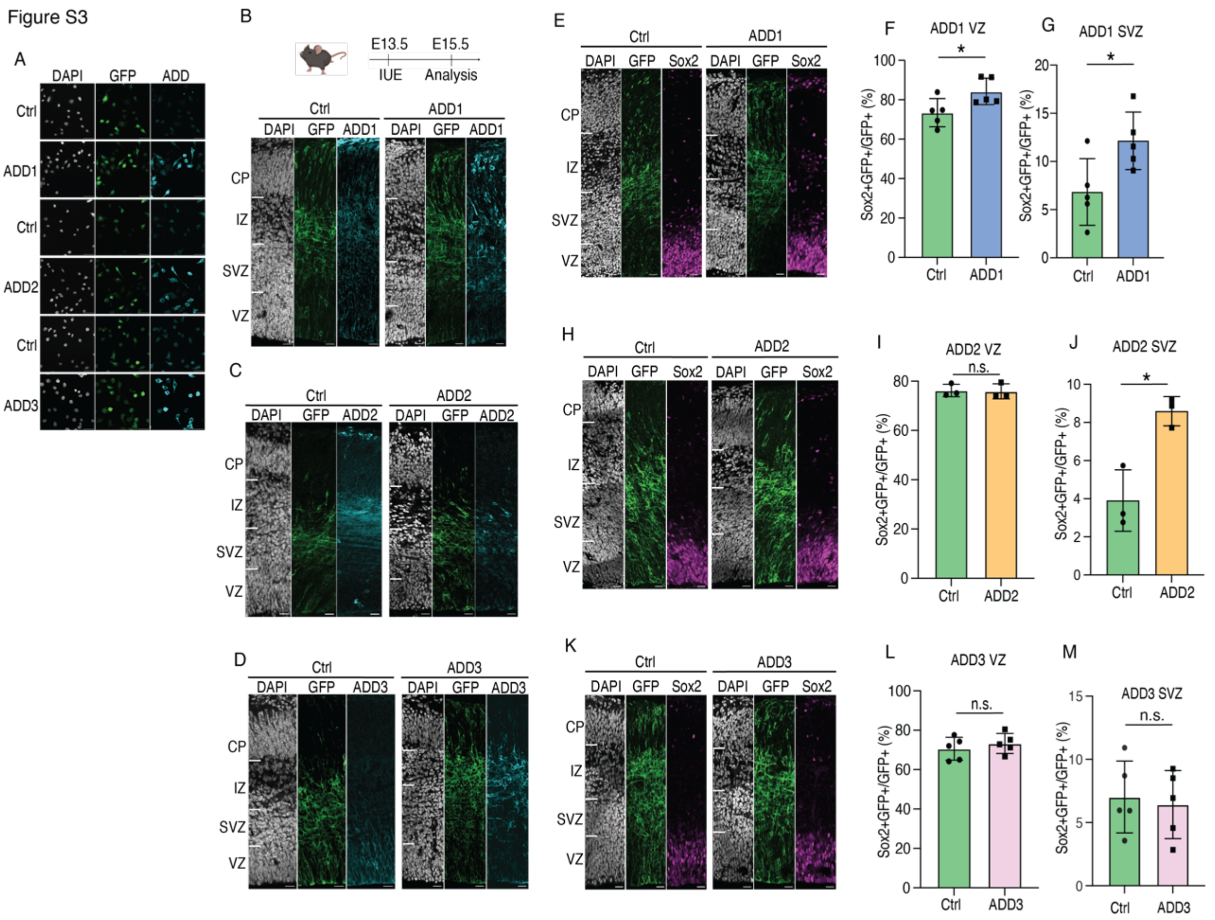


**Figure S3. ADD1 and ADD2, but not ADD3 promote BP abundance in mouse embryonic neocortex.**

(A) ADDs overexpression was tested in HeLa cells. Cells were transfected with empty pCAGGs plasmid as Control (Ctrl) or ADD1, ADD2 or ADD3 along with GFP and analyzed 24 h later by DAPI staining and IF for GFP and ADD1, ADD2, ADD3 respectively (indicated ADD on the panel). Scale bar, 50 μm.

(B-M) ADDs were electroporated along with LynGFP (revealing cell membrane) overexpressing plasmid at E13.5 in mouse embryonic neocortex and embryos were analyzed at E15.5.

(B-D) DAPI staining and IF for GFP and ADD1 (B), ADD2 (C) and ADD3 (D) for control (left) and ADD OE (right). Note that ADDs overexpression increases ADDs protein levels. Scale bars, 20 μm.

(E, H, K) DAPI staining and IF for GFP and Sox2 for control (left) and ADD1 (E), ADD2 (H) and ADD3 (K) (right). Scale bars, 20 µm

(F, I, L) OE of ADD1 (F), but not ADD2 (I) and ADD3 (L), increases Sox2+ APs in mouse developing neocortex. N= 3-5 control and 3-5 OE embryos from 3 different litters. *$p < 0.05$, n.s. not statistically significant; Student's *t* test.

(G, J, M) OE of ADD1 (G) and ADD2 (J), but not ADD3 (M), increases Sox2+ BPs in mouse developing neocortex. N=5 control and 5 OE embryos (G, M) or 3 control and 3 OE embryos (J) from 3 different litters. *$p < 0.05$, n.s. not statistically significant; Student's t test.

Figure S4

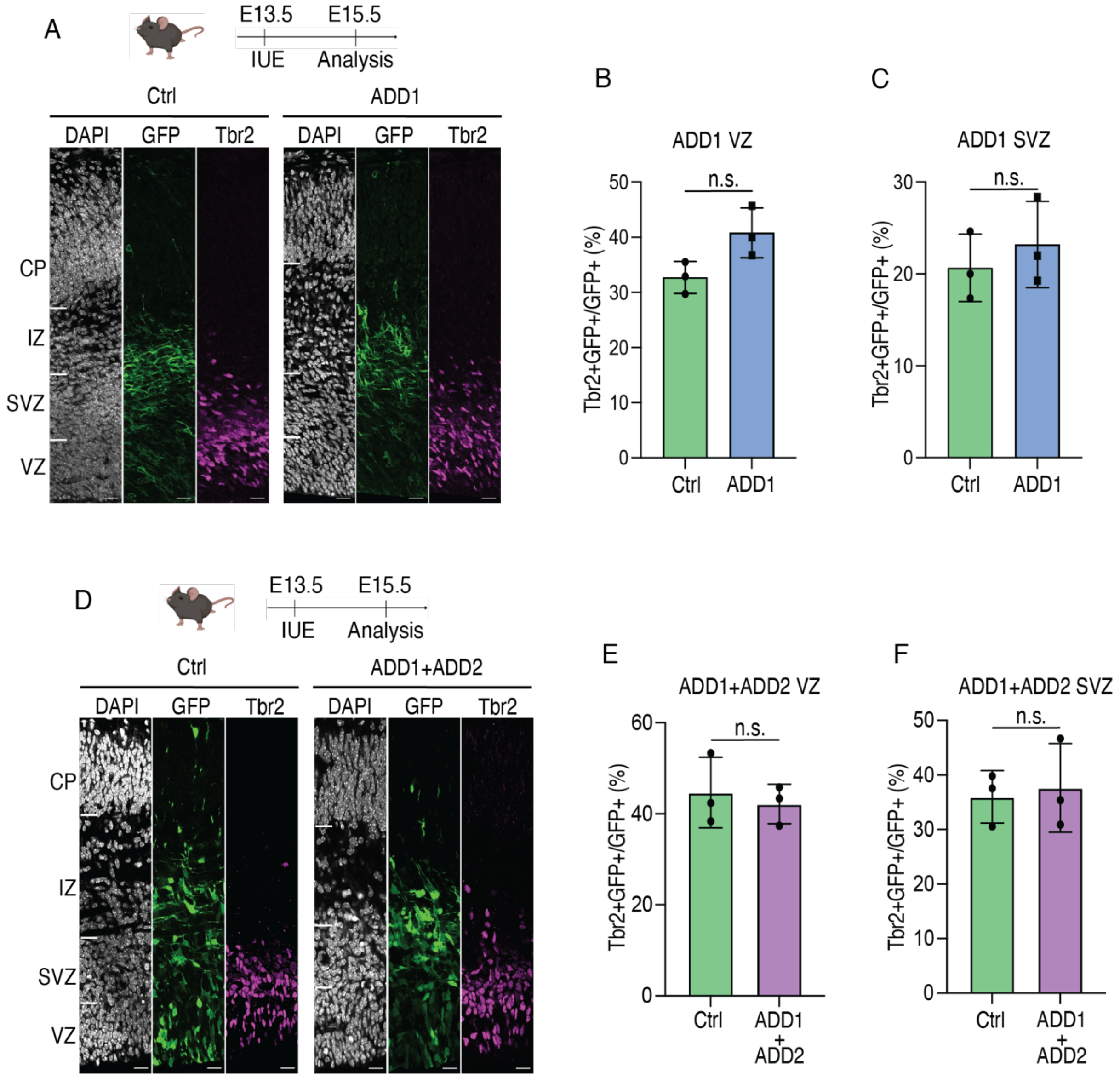


**Figure S4. ADDs overexpression does not affect abundance of Tbr2+ NPCs.**

ADDs OE in mouse developing neocortex does not affect Tbr2+ progenitors' abundance in VZ and SVZ. IUE was performed at E13.5 and analysis at E15.5.

(A, D) DAPI staining and IF for GFP and Tbr2 for control (A and D, left) and ADD1OE (A, right), and ADD 1 and ADD2 co-OE (D, right). Scale bar 20 µm.

(B, C, E, F) ADD1 OE (B, C) and ADD1 and ADD2 co-OE (E, F) do not affect Tbr2+ NPCs in VZ (B, E) and SVZ (C, F). N=3 control and 3 OE/co-OE embryos from 3 different litters. n.s. not statistically significant, Student's *t* test.

Figure S5

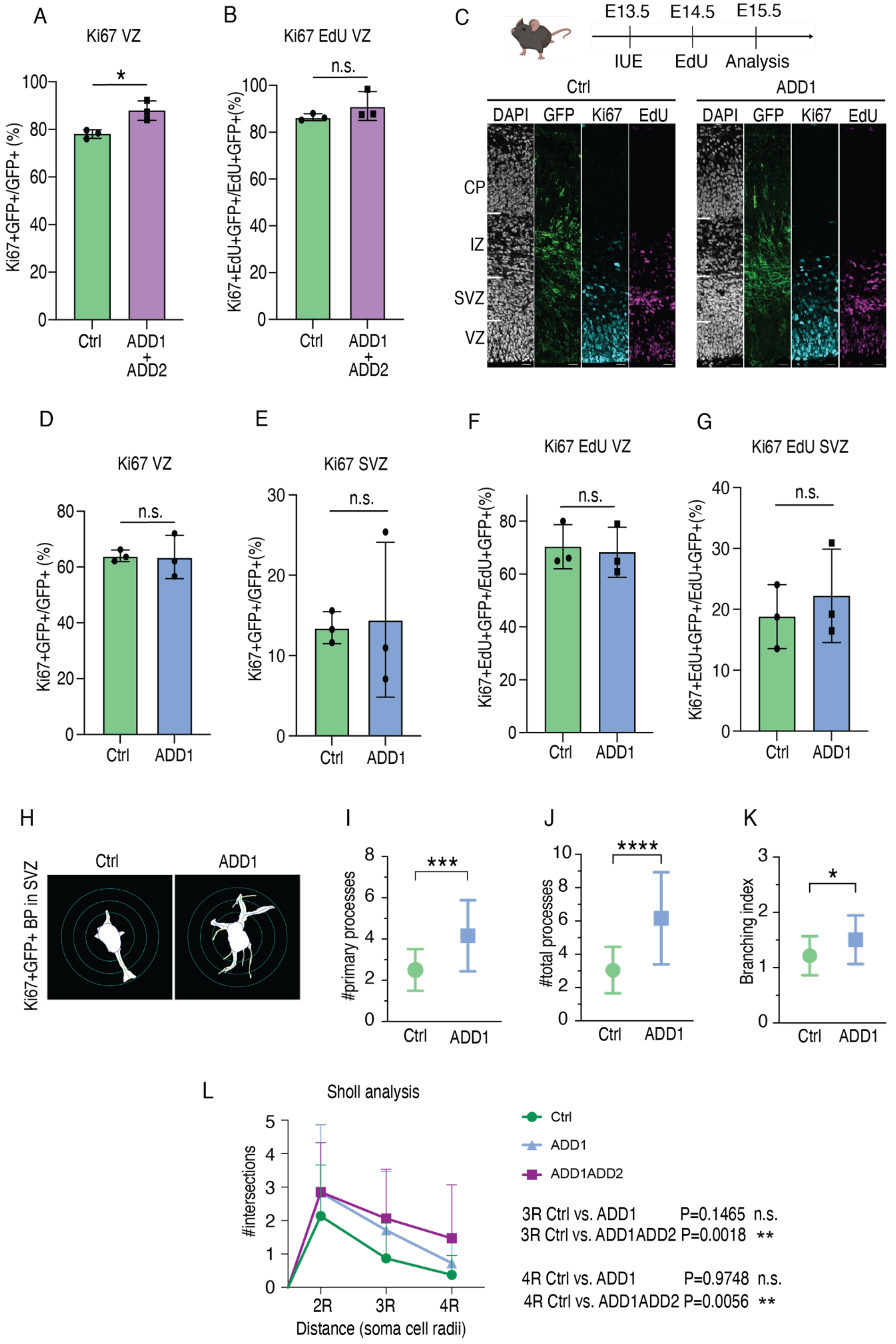

A
Ki67 VZ
*
Ki67+GFP+/GFP+ (%)
Ctrl
ADD1 + ADD2
B
Ki67 EdU VZ
n.s.
Ki67+EdU+GFP+/EdU+GFP+(%)
Ctrl
ADD1 + ADD2
C
E13.5
E14.5
E15.5
IUE
EdU
Analysis
Ctrl
ADD1
DAPI
GFP
Ki67
EdU
CP
IZ
SVZ
VZ
D
Ki67 VZ
n.s.
Ki67+GFP+/GFP+ (%)
Ctrl
ADD1
E
Ki67 SVZ
n.s.
Ki67+GFP+/GFP+(%)
Ctrl
ADD1
F
Ki67 EdU VZ
n.s.
Ki67+EdU+GFP+/EdU+GFP+(%)
Ctrl
ADD1
G
Ki67 EdU SVZ
n.s.
Ki67+EdU+GFP+/EdU+GFP+(%)
Ctrl
ADD1
H
Ki67+GFP+ BP in SVZ
Ctrl
ADD1
I
***
#primary processes
Ctrl
ADD1
J
****
#total processes
Ctrl
ADD1
K
*
Branching index
Ctrl
ADD1
L
Sholl analysis
#intersections
2R
3R
4R
Distance (soma cell radii)
Ctrl
ADD1
ADD1ADD2
3R Ctrl vs. ADD1 P=0.1465 n.s.
3R Ctrl vs. ADD1ADD2 P=0.0018 **
4R Ctrl vs. ADD1 P=0.9748 n.s.
4R Ctrl vs. ADD1ADD2 P=0.0056 **

**Figure S5. BPs morphology and proliferation upon ADDs OE.**

(A, B) ADD1 and ADD2 co-OE increases NPC abundance but not proliferative capacity in the VZ. ADDs were electroporated along with lynGFP overexpressing plasmid at E13.5 in mouse embryonic neocortex, EdU was administered by IP injection at E14.5 and embryos were analyzed at E15.5. ADD1 and ADD2 co-OE increases Ki67+ actively proliferating APs (A), but not cell cycle re-entry (B). N=3 control and 3 co-OE embryos from 3 different litters. $^*p < 0.05$, n.s. not statistically significant; Student's *t* test.

(C-G) ADD1 does not increase NPC proliferative capacity or cell cycle re-entry. ADD1 was electroporated along with lynGFP overexpressing plasmid at E13.5 in mouse embryonic neocortex, EdU was administered by IP injection at E14.5 and embryos were analyzed at E15.5.

(C) DAPI staining and IF for GFP, Ki67 and EdU of E15.5 mouse neocortex for control (left) and ADD1 OE (right). Scale bar 20 µm.

(D-G) ADD1 OE does not affect abundance of Ki67+ actively proliferating APs in the VZ (D) and BPs in the SVZ (E), nor does it affect the cell cyle re-entry of NPC in VZ (F) and SVZ (G). N= 3 control and 3 OE embryos from 3 different litters. n.s. not statistically significant, Student's *t* test.

(H-K) ADD1 OE increases the abundance of BP protrusions in mouse developing neocortex. ADD1 was electroporated along with GFP overexpressing plasmid at E13.5 in mouse embryonic neocortex and analyzed at E15.5.

(H) Segmentation masks of Ki67+GFP+ BPs in mouse E15.5 SVZ of control (left) and ADD1 OE (right).

(I-K) ADD1 OE increases the number of BPs primary (I) and total (J) processes as well as the branching index (K).

N=22 control and 25 ADD1 OE cells from 3 different litters. $^{****}p < 0.001$, $^{***}p < 0.001$; $^*p < 0.05$; Mann-Whitney u-test (I, K); Welch's t test (J).

(L) Sholl analysis of BPs. ADD1 and ADD2 co-OE, but not ADD1 OE, increases the abundance of longer BP protrusions (3R and 4R). ADDs were electroporated along with GFP overexpressing plasmid at E13.5 in mouse embryonic neocortex and analyzed at E15.5. N=45 control, 25 ADD1 OE and 34 ADD1 and ADD2 co-OE cells from 3 different litters. $^{**}p < 0.01$; Sidak's multiple comparisons test, 2way ANOVA.

Figure S6

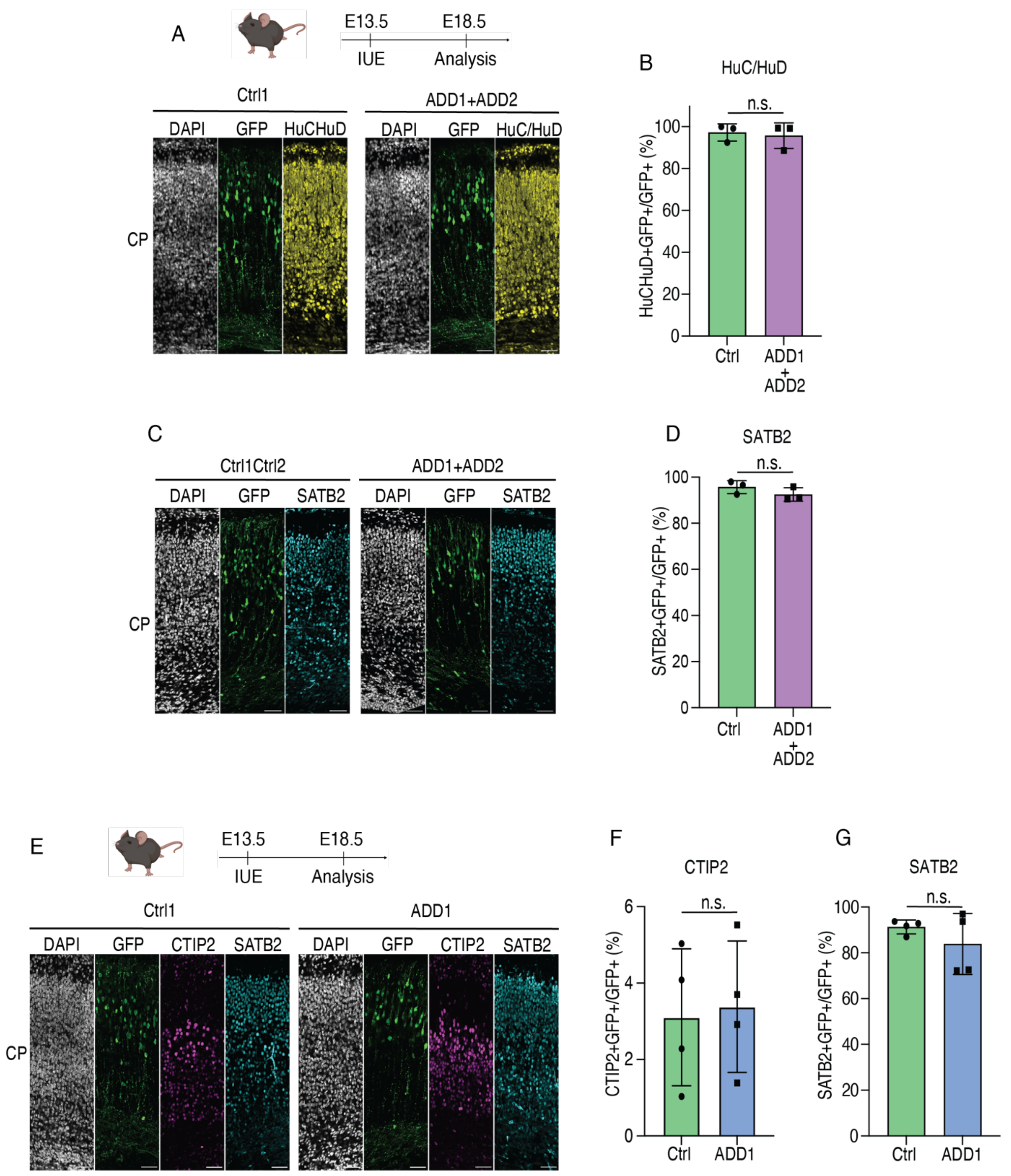


**Figure S6. Effects of ADDs OE on neurogenesis at late stages of development.**

(A-D) ADD1 and ADD2 were co-electroporated along with GFP overexpressing plasmid at E13.5 in mouse embryonic neocortex and embryos were analyzed at day E18.5. (A, C) DAPI staining and IF for GFP and HuC/HuD (A) or Satb2 (C) of E18.5 mouse neocortex for control (left) and ADD1 and ADD2 co-OE (right). Scale bars, 50 μm.

(B, D) Vast majority of GFP+ cells in both control and co-OE at E18.5 are HuC/HuD+ (B) and Satb2+ (D) neurons. N= 3 control and 3 co-OE embryos from 3 different litters. n.s. not statistically significant; Student's *t* test.

(E-G) ADD1 was electroporated along with GFP overexpressing plasmid at E13.5 in mouse embryonic neocortex and embryos were analyzed at day E18.5.

(E) DAPI staining and IF for GFP, CTIP2 and SATB2 of E18.5 mouse neocortex for control (left) and ADD1 (right). Scale bars, 50 μm.

(F, G) ADD1 OE does not affect abundance of Ctip2+(F) and Satb2+ (G) neurons. N=4 control and 4 OE embryos from 3 different litters. n.s. not statistically significant, Student's *t* test.

Figure S7

A

| ADD1 | | | | off-targets | | | | |
|---|---|---|---|---|---|---|---|---|
| | SPACER | % pred.eff. | exon | MM0 | MM1 | MM2 | MM3 | MM4 |
| sg1 | GATCCTCTAAGATCGTTCAC | 60,49 | 4 | 0 | 0 | 0 | 0 | 24 |
| sg2 | GACCAGAGTGAACTCCGAGC | 59,89 | 3 | 0 | 0 | 0 | 3 | 66 |
| sg3 | TTTGGAAAAGGTTCGAACTC | 51,68 | 7 | 0 | 0 | 0 | 1 | 88 |

B

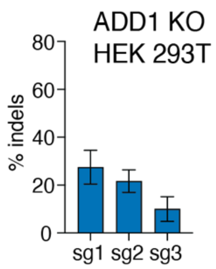


C Indels per H9 clone

| Clone ID | Allele 1 | Allele 2 |
|---|---|---|
| 1_C11 | -5 | -1 |
| 1_E7 | -25 | -1 |
| 2_B12 | -14 | -1 |

D Chromatograms per H9 clone

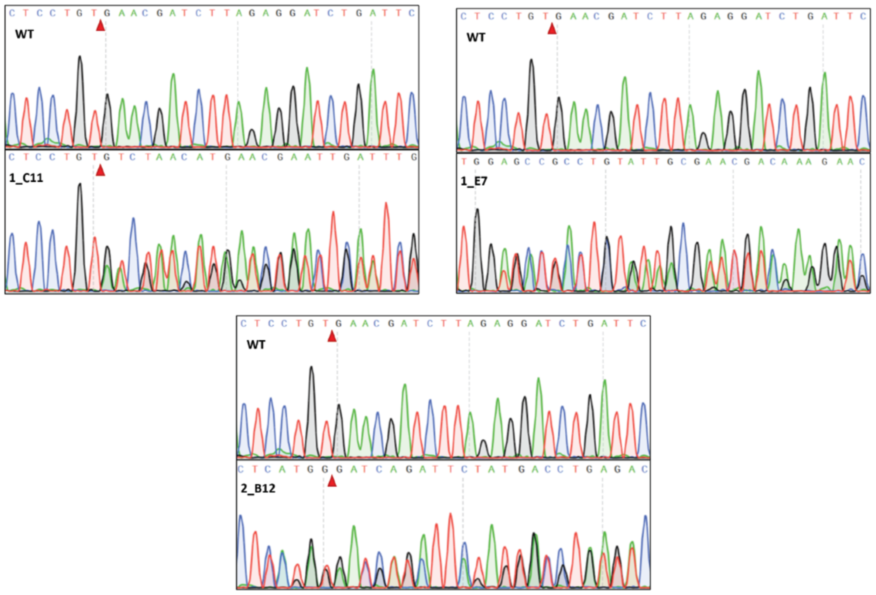


**Figure S7. Generation of ADD1 KO pluripotent stem cells (PSC)**

A) Characteristics of the single guide (sg) RNAs used for the editing of the ADD1 locus in HEK293T cells. The name and sequence of the sgRNAs is reported along with the targeted exon and the percentage of in-silico predicted efficiency (% pred.eff.) using the CHOPCHOP web tool. Off-Targets; the numbers refer to the predicted off-targets with increasing Mismatches (MM) from the protospacer sequence (MM0, MM1, MM2, MM3, MM4).

B) Validation of sgRNAs in HEK293T cells. The efficiency of each sgRNA is measured as percentage of indels, normalized on a scrambled control sgRNA (sgCTRL). Data are means of 3 (for sg2 and sg3) or 4 (for sg1) biological replicates. Error bars indicate SD. Note that the gene editing in the H9 PSCs was done using the sg1.

C) Characterization of the three H9 PSC clones carrying mutations in ADD1 locus obtained upon gene editing with the sg1. Indels are deconvoluted for each allele using the Synthego's ICE software. Note that the data show that in all three clones both alleles have a frameshift deletion.

D) Sanger chromatograms of WT H9 cells and the three KO clones. Red triangle, the cutting site. Note that for the clone 1E7 the absence of the red triangle in the edited clone is due to the indel being too large and it disrupted the protospacer sequence.

Figure S8

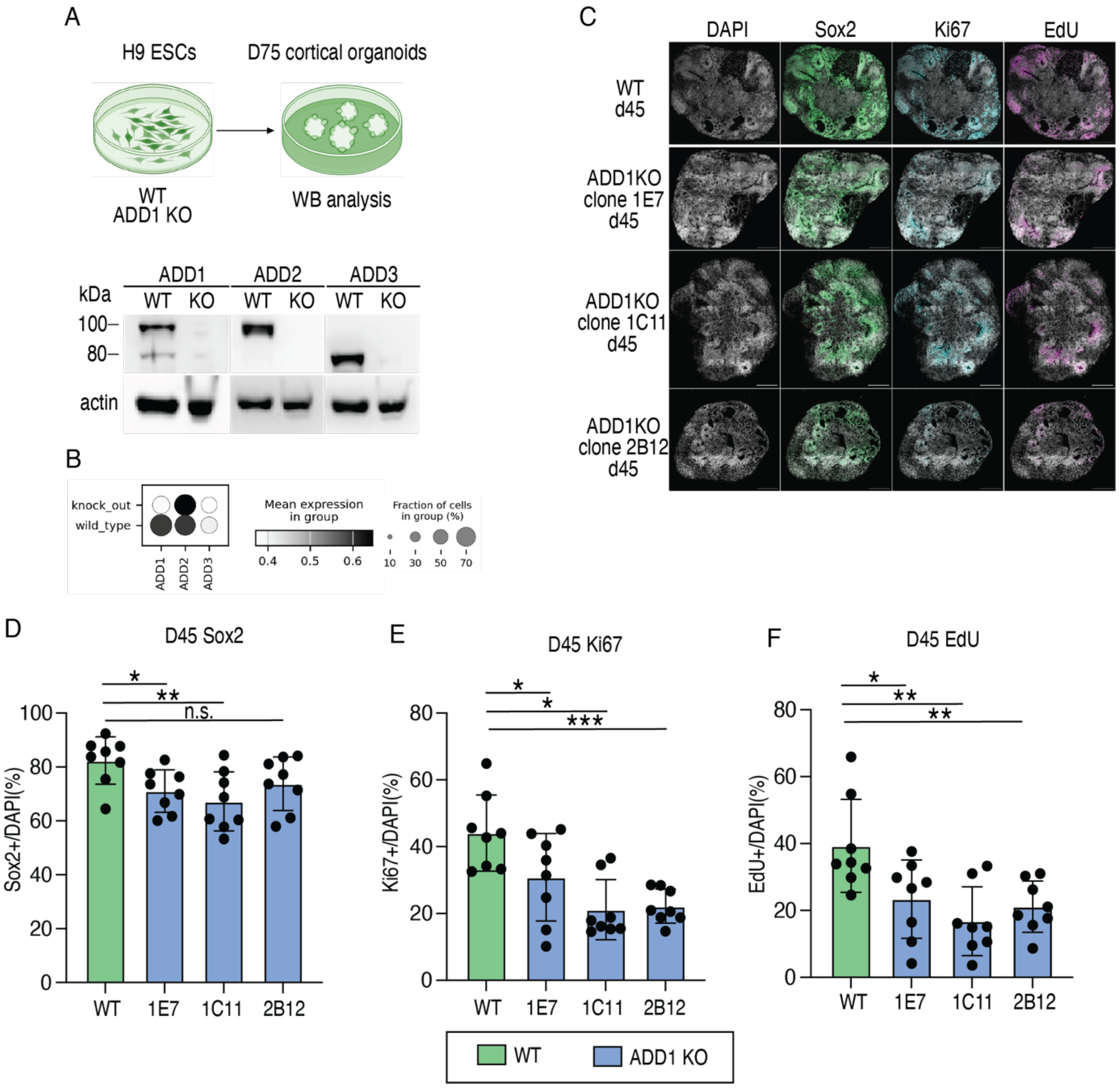


**Figure S8. ADD1 KO in reduces NPC proliferation in human cortical organoids**

Cortical organoids derived from WT and ADD1 KO cells were obtained as in Figure 4 and analyzed at day 45 (C-F) or 75 (A, B).

(A) Immunoblot for ADD1 (left), ADD2 (middle), ADD3 (right) and actin (down) showing absence of all three ADDs upon ADD1 KO.

(B) Mean expression of ADD1, ADD2 and ADD3 mRNA as detected by scRNA-seq (see Figure 5), showing lack of *ADD1*, but presence of *ADD2* and *ADD3* transcripts.

(C) DAPI staining and IF for Sox2, Ki67 and EdU of WT (upper) and three ADD1 KO clones (1E7, 1C11, 2B12) in day 45 cortical organoids. Scale bar 100 µm.

(D-F) ADD1 KO leads to a reduction in the abundance of SOX2+ (D) and Ki67+ (E) NPCs and EdU+ proliferative cells (F) in day 45 cortical organoids. N = 8 ventricles from 3 different differentiation batches. ***$p < 0.001$, **$p < 0.01$, *$p < 0.05$, n.s. not statistically significant; Welch's *t* test.

Figure S9

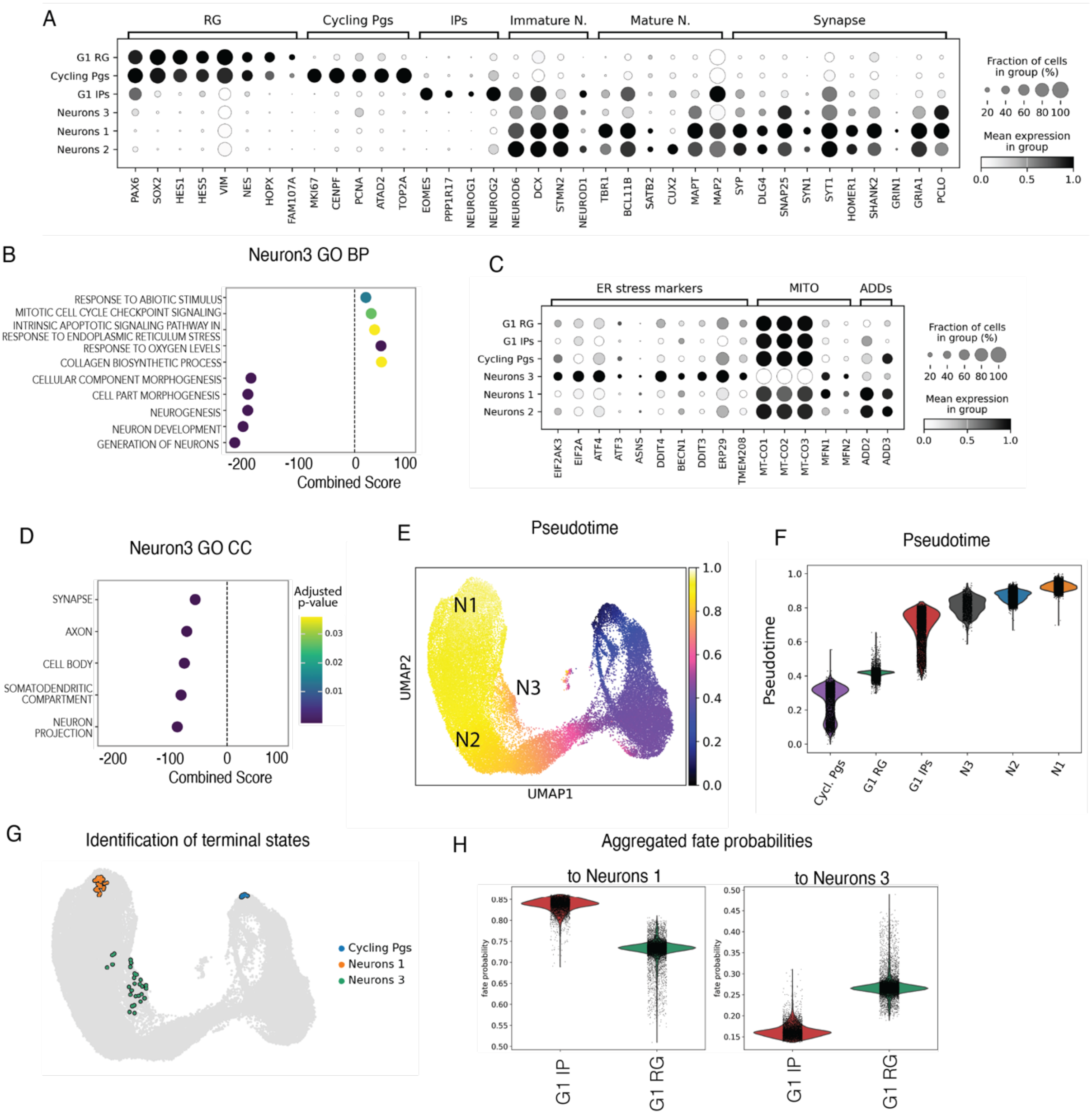


**Figure S9. Neurons 3 are enriched in ADD1 KO, they exhibit signature of ER stress, aberrant neurogenesis and are likely generated by RG.**

ScRNA-seq analysis of WT and ADD1 KO cortical organoids as described in Figure 5

(A) Dotplot showing markers used for annotation of the principal six cell clusters. Note, that the G1 RG and Cycling progenitors clusters both contain apical and basal RG (the latter revealed with positivity for HOPX and FAM107A). Whereas canonical markers of immature neurons are equally expressed in both Neurons 1 and 2, we detected a higher mean expression of various synaptic genes in Neurons 1 compared to Neurons 2, suggesting their more advanced maturity. We further detected a strong enrichment of markers for lower cortical layers (TBR1 and CTIP2) in Neurons 1 and similar enrichment of markers for upper cortical layers (CUX2) in Neurons 2. As to Neurons 3, they showed reduced expression of overall neuronal and synaptic markers, apart from PCLO, which is however known to exhibit also non-synaptic roles (Ahmed et al., 2015).

(B, D) Plot showing gene ontology Biological process (B) and Cellular component (D) terms enriched in Neurons 3 compared to Neurons 2. Note the up-regulation of genes associated with the ER stress and the down-regulation of genes linked to neurogenesis progression and cell morphology.

(C) Dotplot showing higher mean expression of markers associated to ER stress and reduced mean expression of mitochondrial genes in the Neuron 3 cluster. In addition, this cluster also exhibited reduced mean expression of ADD2 and ADD3 compared to the other two neuronal clusters.

(E) Reconstruction of the pseudotime trajectory. The cell with the highest TOP2A (in the cycling progenitors cluster) expression served as the root cell of the trajectory inference. N1-N3, neuronal clusters 1-3.

(F) Violin plot of the distribution of the main clusters along the pseudotime trajectory. Note that Neurons 3 appear before the other neuronal clusters on the pseudotime trajectory.

(G) Terminal states recovered by CellRank. Terminal states corresponding to Cycling progenitors and Neurons 1 were recovered automatically. Terminal state corresponding to Neurons 3 was recovered by selecting the 30 cells with highest PERK expression.

(H) Aggregated fate probabilities calculated by CellRank showing fate probability of IPs and RG towards either of terminal states (Neurons 1, left; Neurons 3, right). Note that aberrant Neurons 3 are more likely generated by RG than IPs.

Figure S10

A

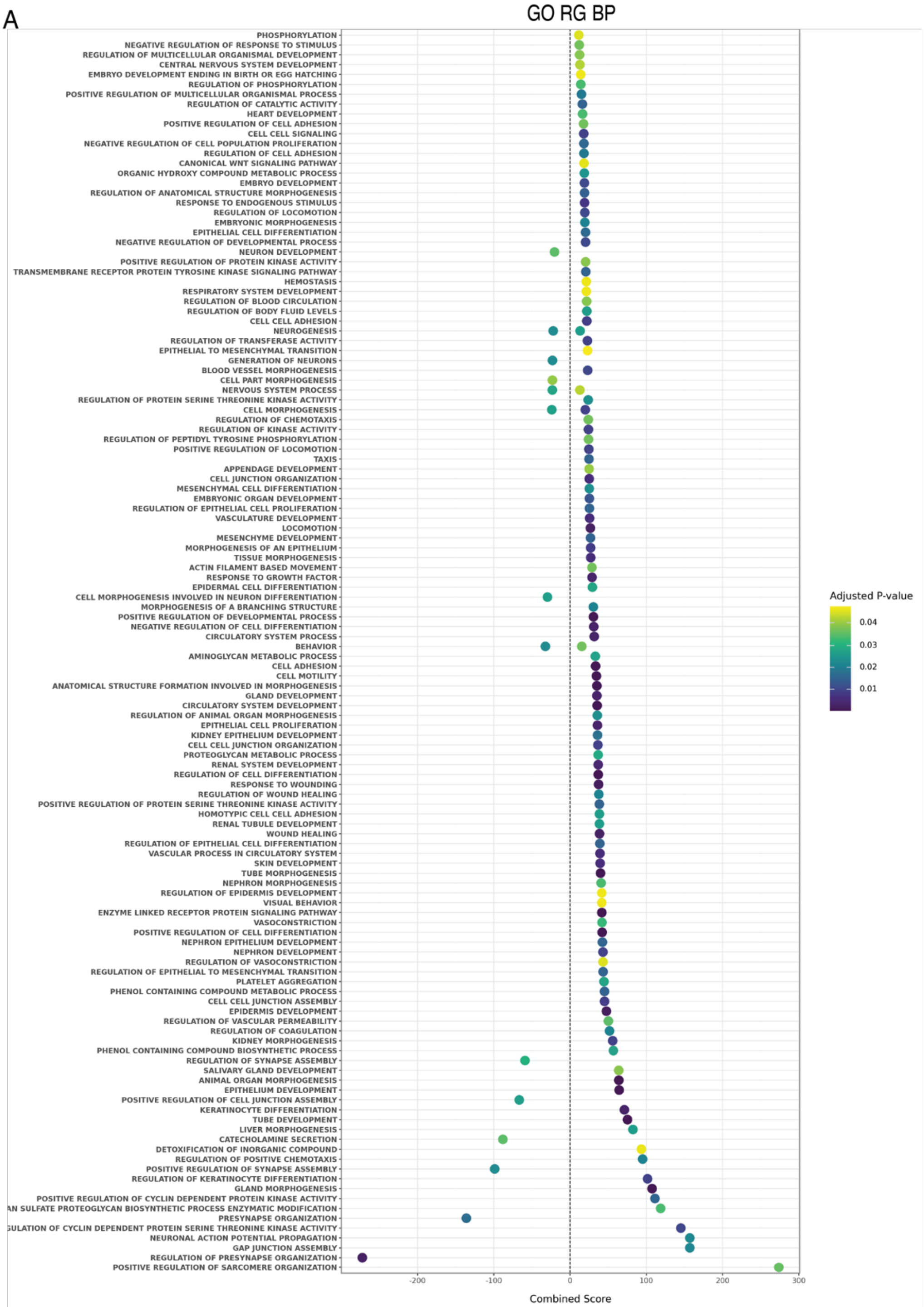


B

Down-regulated in neurogenesis

SLC4A10, EFNB1, SLITRK5, OLFM3, PCSK9, HDAC9, CDHR1, SLITRK1, POTEF, FOXO6, FLRT2,CDH23, UNC5D, WNT16, SLITRK6, WNT7A, NRP1, SH3GL2, DMRT3, TNR, EML1, IL1RAPL1, SPAG6, POU3F1, SYT1, FSTL4, NFASC, PLAG1, CDKN2C, KCNQ3, PRDM8, ATP2B2, RNF220, MCF2

Up-regulated in neurogenesis

EPB41L3, SIPA1L1, PLXNA4, CXCL12, SDC2, SERPINE2, EPOP, IRX1, ZEB2, GFAP, DCT, IRX5, DCLK1, FZD5, CEBPB, ANOS1, SMARCA1, TNC, MYO16, UST, EGFR, PROX1, NBL1, DAAM2, S100B, ECE1, TGFB2, C1QL1, GPR37, SCN1B, ADGRG6, ZIC3, CLCN2, NKD1, EFNA1, EDNRB, MYLIP, LYN, NPY, CSF1, OLIG2, NEURL1, NRCAM, NTF3, IRX2

Down-regulated in cell morphogenesis

EFNB1, SLITRK5, CDHR1, SLITRK1, POTEF, FLRT2, GAS2, CDH23, UNC5D, SLITRK6, WNT7A, NRP1, SH3GL2, DMRT1, TNR, IL1RAPL1, SPAG6, SYT1, FSTL4, CDH7, NFASC, PRDM8, MCF2

Up-regulated in cell morphogenesis

EPB41L3, SIPA1L1, RHOJ, PLXNA4, CXCL12, COL18A1, SDC2, MYH9, HPN, ACTN1, CDH20, DCLK1, ANOS1, LRATD1, MYO16, EGFR, PROX1, NBL1, S100B, ECE1, TGFB2, FERMT2, LATS2, SCN1B, CD44, EFNA1, CDH13, BAMBI, MYH14, NRCAM, NTF3, UST

**Figure S10. GO terms enriched in RG upon ADD1 KO.**

ScRNA-seq analysis of WT and ADD1 KO cortical organoids as described in Figure 5

(A) Plot showing the complete list of gene ontology (biological process) terms enriched in radial glia (GO RG BP) in ADD1 KO vs. WT.

(B) Lists of genes contributing to the gene ontologies “Neurogenesis” and “Cell Morphogenesis” which were both up- and down-regulated upon ADD1 KO. Note that both Neurogenesis- and Cell Morphogenesis associated genes down-regulated upon ADD1 KO gene prominently belong to guidance molecules and their receptors along with the cell adhesion molecules, whereas Neurogenesis-associated genes up-regulated upon ADD1 KO are associated with cell migration, gliogenesis and negative regulation of the cell cycle progression and cell adhesion.

Figure S11

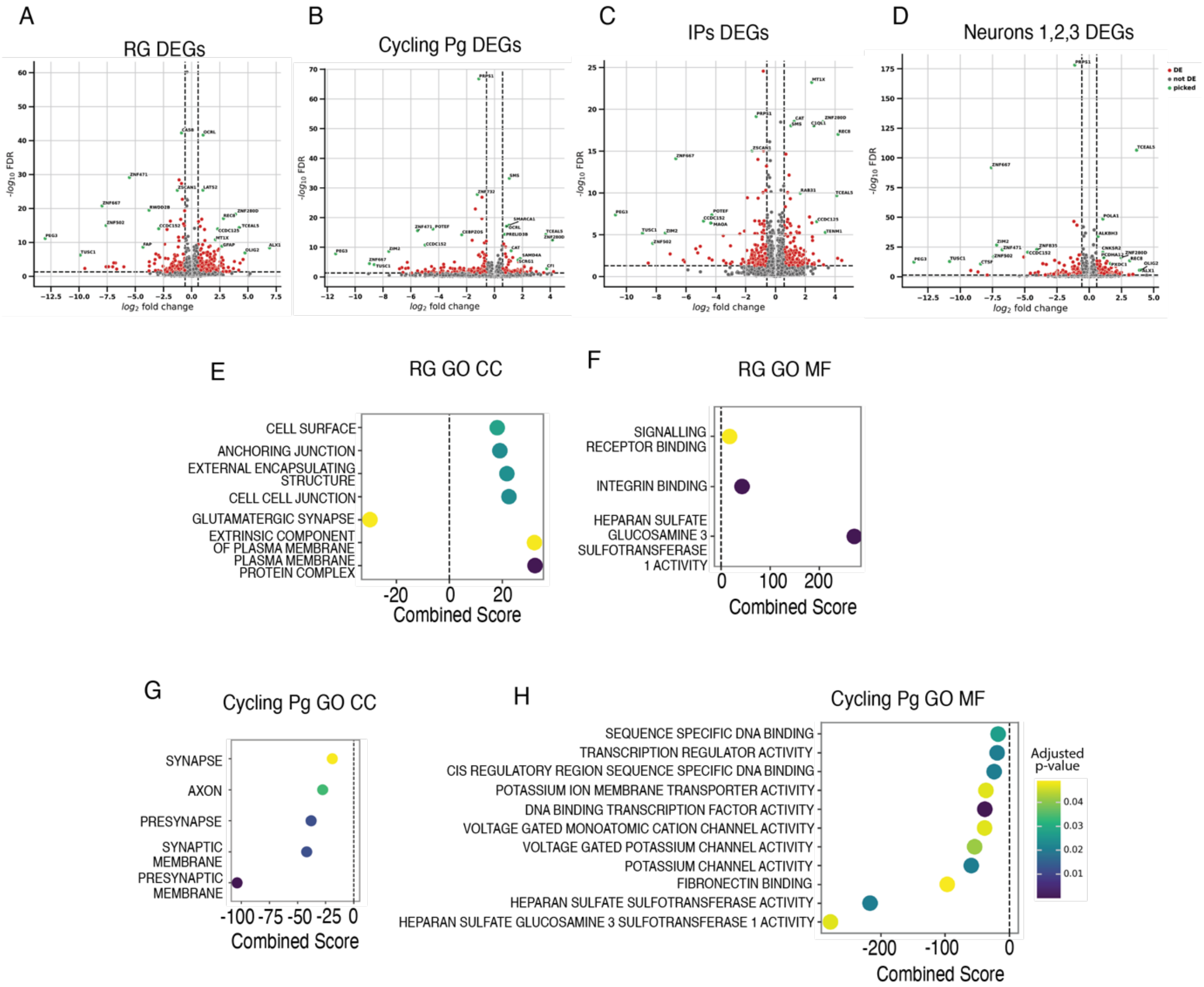


**Figure S11. GO terms enriched upon ADD1 KO.**

ScRNA-seq analysis of WT and ADD1 KO cortical organoids as described in Figure 5

(A-D) Volcano plots showing differentially expressed genes (red) in radial glia (A), cycling progenitors (B), intermediate progenitors (C) and neurons (D, combined clusters Neurons 1, 2 and 3) upon ADD1 KO. Top genes are indicated in green.

(E-H) Plots showing gene ontologies Cellular component (E, G) and Molecular function (F, H) terms enriched in Radial glia (E, F) and Cycling progenitors (G, H), upon ADD1 KO.

**Table S1. List of oligonucleotides used**

| Oligonucleotide | Sequence |
|---|---|
| ADDs cloning primers | |
| ADD1 forward | CGATTCGCTTCTGAGGAACCT |
| ADD1 reverse | ACAAGGACAGAGCACAGACG |
| ADD2 forward XhoI | TATACTCGAGCACCGGGAAAATGAGC |
| ADD2 reverse Xho I | TATACTCGAGTGCAGGGACAGAGATG |
| ADD3 forward XhoI | TATCCTCGAGGAGTAATCCACAGAC |
| ADD3 reverse BglII | TATAAGATCTTTAGGCCTCAACT |
| ADD1 guides amplifying primers | |
| sg1 ADD1 Fw | GTCTCGCTTTGTTGCCTTGG |
| sg1 ADD1 Rev | CTTACTGAAGCTCAGCCTGC |
| sg2 ADD1 Fw | CCTGTCAGAAGGCTAGTGTG |
| sg2 ADD1 Rev | CCAGGCTGGTCTCGAACTTA |
| sg3 ADD1 Fw | GTCTGTGGCGTAGTGAAGTC |
| sg3 ADD1 Rev | GACAGCAATCGCACTTGCTG |

**References cited in the Supplemental figure legends:**

Ahmed, M.Y., Chioza, B.A., Rajab, A., Schmitz-Abe, K., Al-Khayat, A., Al-Turki, S., Baple, E.L., Patton, M.A., Al-Memar, A.Y., Hurles, M.E.*, et al.* (2015). Loss of PCLO function underlies pontocerebellar hypoplasia type III. Neurology *84*, 1745-1750.

Fietz, S.A., Lachmann, R., Brandl, H., Kircher, M., Samusik, N., Schroder, R., Lakshmanaperumal, N., Henry, I., Vogt, J., Riehn, A.*, et al.* (2012). Transcriptomes of germinal zones of human and mouse fetal neocortex suggest a role of extracellular matrix in progenitor self-renewal. Proceedings of the National Academy of Sciences of the United States of America *109*, 11836-11841.

Florio, M., Albert, M., Taverna, E., Namba, T., Brandl, H., Lewitus, E., Haffner, C., Sykes, A., Wong, F.K., Peters, J.*, et al.* (2015). Human-specific gene ARHGAP11B promotes basal progenitor amplification and neocortex expansion. Science *347*, 1465-1470.

Qian, X., Su, Y., Adam, C.D., Deutschmann, A.U., Pather, S.R., Goldberg, E.M., Su, K., Li, S., Lu, L., Jacob, F.*, et al.* (2020). Sliced Human Cortical Organoids for Modeling Distinct Cortical Layer Formation. Cell Stem Cell *26*, 766-781 e769.

(A)
